# Inhibitors of ERp44, PDIA1, and AGR2 induce disulfide-mediated oligomerization of Death Receptors 4 and 5 and cancer cell death

**DOI:** 10.1101/2021.01.13.426390

**Authors:** Mary E. Law, Elham Yaaghubi, Amanda F. Ghilardi, Bradley J. Davis, Renan B. Ferreira, Jin Koh, Sixue Chen, Sadie F. DePeter, Christopher M. Schilson, Chi-Wu Chiang, Coy D. Heldermon, Peter Nørgaard, Ronald K. Castellano, Brian K. Law

## Abstract

Breast cancer mortality remains unacceptably high, indicating a need for safer and more effective therapeutic agents. Disulfide bond Disrupting Agents (DDAs) were previously identified as a novel class of anticancer compounds that selectively kill cancers that overexpress the Epidermal Growth Factor Receptor (EGFR) or its family member HER2. DDAs kill EGFR+ and HER2+ cancer cells via the parallel downregulation of EGFR, HER2, and HER3 and activation/oligomerization of Death Receptors 4 and 5 (DR4/5). However, the mechanisms by which DDAs mediate these effects are unknown. Affinity purification analyses employing biotinylated-DDAs reveal that the Protein Disulfide Isomerase (PDI) family members AGR2, PDIA1, and ERp44 are DDA target proteins. Further analyses demonstrate that shRNA-mediated knockdown of AGR2 and ERp44, or expression of ERp44 mutants, enhance basal and DDA-induced DR5 oligomerization. DDA treatment of breast cancer cells disrupts PDIA1 and ERp44 mixed disulfide bonds with their client proteins. Together, the results herein reveal DDAs as the first small molecule, active site inhibitors of AGR2 and ERp44, and demonstrate roles for AGR2 and ERp44 in regulating the activity, stability, and localization of DR4 and DR5, and activation of Caspase 8.

## Introduction

In spite of significant improvements in early detection, and new molecularly targeted therapeutics, breast cancer remains the second largest cancer killer of women in America after lung cancer, and is the most frequently diagnosed cancer in American women [1]. Much of the lethality of breast cancer derives from our inability to successfully treat patients with metastatic and/or drug-resistant malignancies. New drugs are needed that bypass resistance to currently used regimens by targeting novel cancer cell-specific vulnerabilities. A class of agents termed Disulfide bond Disrupting Agents (DDAs) was previously identified and shown to kill breast cancer cells in association with downregulation of the HER-family proteins EGFR, HER2, and HER3, and decreased activating phosphorylation of the Akt kinase [2]. Subsequent work showed that DDAs induce endoplasmic reticulum stress (ERS) [3] and alter the pattern of disulfide bonding of EGFR and the TRAIL receptors, Death Receptor 4 (DR4) and Death Receptor 5 (DR5) [4]. DDA-induced cell death is mediated by activation of the Caspase 8-Caspase 3 extrinsic apoptotic pathway.

Little is known regarding how the disulfide bonding of HER-family and Death Receptor-family proteins are chaperoned within the endoplasmic reticulum, and how the disulfide bonding patterns of these proteins control their stability, intracellular localization, and downstream signaling. TNF-Related Apoptosis Inducing Ligand (TRAIL) has been considered a promising cancer therapeutic agent due to its selectivity for killing cancer cells, while leaving normal tissues unharmed [5–8]. However, TRAIL analogs and DR4/5 agonistic antibodies have yet to be clinically approved for anticancer therapy. This is due in part to pharmacokinetic issues related to these protein drugs themselves, and in part to the ability of cancer cells to acquire resistance through a variety of mechanisms, including downregulation of DR4 and DR5 (reviewed in [9]). Since DDAs upregulate DR5 and activate DR4 and DR5 in a ligand-independent manner [4, 10], DDAs may overcome the resistance mechanisms that suppress the efficacy of other activators of the TRAIL/Death Receptor pathway. A recent report showed that the extracellular domain of DR5 prevents oligomerization and downstream activation of Caspase 8 in the absence of ligand engagement, and that excision of the DR5 extracellular domain permits full activation of DR5 in a ligand-independent manner [11]. It was previously proposed that DDAs activate DR5 in a ligand-independent manner by permitting inter-molecular disulfide bond formation at the expense of intramolecular disulfide bonding, resulting in DR5 oligomerization and activation [4, 10]. However, the precise mechanisms by which this happens are unknown. Similarly, DDA treatment causes disulfide bond-dependent EGFR oligomerization followed by degradation [2, 4], but how DDAs perturb EGFR disulfide bonding is unknown. The Protein Disulfide Isomerase (PDI) family member AGR2 is essential for EGFR folding and surface localization [12], and likewise facilitates expression of Mucins [13, 14].

The PDI family contains 21 members (reviewed in [15]). Canonical PDIs are considered promising targets for anticancer therapeutics [16] and significant attention has focused on the roles of canonical PDIs in mediating disulfide bond oxidation, reduction, or exchange via their active site thioredoxin-like CXXC motifs. However, other PDI family members are less well studied. The PDIs AGR2 and AGR3 contain a non-canonical CXXS active site motif. Based on biochemical PDI mutagenesis studies [17] it is predicted that AGR2/3 will exhibit disulfide isomerase activity, but not oxidase or reductase activity. Consistent with this, AGR2 forms stable mixed disulfide bonds with its client proteins [12, 13]. Another PDI family member, ERp44, also contains an active site CXXS motif, and has multiple functions within the endoplasmic reticulum (ER). ERp44 plays a critical role in the disulfide-mediated oligomerization of Adiponectin [18, 19] and Immunoglobulin M [20, 21]. Additionally, ERp44 anchors client/partner proteins such as ERO1α, Prx4, FGE/SUMF1, and ER Aminopeptidase 1 (ERAP1) [22–25] that do not possess C-terminal -KDEL ER retention sequences to prevent their secretion (reviewed in [26]).

HER-family proteins are the only sub-family of receptor tyrosine kinases that contain two cysteine-rich repeats in their extracellular domain [27]. DR4 and DR5 also harbor numerous disulfide bonds in their extracellular domains [28]. Based on these observations, we previously hypothesized that HER-family proteins and DR4/5 may be particularly sensitive to agents that impede native disulfide bond formation. The mechanisms by which DDAs cause downregulation of EGFR, HER2, and HER3, and activate DR4/5 are unknown. Here we show using novel biotinylated DDA derivatives that DDAs covalently modify ERp44, PDIA1 and AGR2 both in vitro and in intact cells, and that DDA treatment of cells disrupts PDIA1 and ERp44 disulfide bonds with their client proteins, and disulfide-mediated AGR2 dimerization. Together, the results indicate that DDAs activate DR4/5 by inhibiting the activity of the PDI family members ERp44, PDIA1, and AGR2. DDAs are to our knowledge the first active site inhibitors of ERp44 and AGR2.

## Results

### Characterization of novel DDAs with enhanced potency

Previously described DDAs, including RBF3, DTDO, and tcyDTDO (Fig. 1A), were demonstrated to suppress tumor growth through induction of apoptotic cancer cell death [4, 29]. As shown in our previous studies, modifications of the parent cyclic DDA, DTDO, resulted in a more potent DDA, tcyDTDO, with improved pharmacodynamic and pharmacokinetic properties [4, 10, 29]. With tcyDTDO in hand, we then explored the effect of substitutions on its cyclohexane ring to determine if its potency and drug-like properties could be further improved. Our efforts led to the design and synthesis of novel DDAs including dimethoxy-tcyDTDO (dMtcyDTDO) and difluoro-tcyDTDO (dFtcyDTDO) (Fig. 1B). Well appreciated, fluorine’s size and high electronegativity lead to strong C–F bonds that resist metabolic degradation making fluorine substitution an effective way to increase drug stability. For instance, replacing hydrogen with fluorine significantly alters the rate and extent of Cytochrome P450 metabolism [30–33]. Both dMtcyDTDO and dFtcyDTDO exhibit approximately two-fold greater potency than tcyDTDO in cell viability assays (Fig. 1C). DMtcyDTDO also exhibited two-fold greater potency than tcyDTDO in DNA and protein synthesis assays (Fig. S1A). DDAs induce apoptotic cell death associated with ERS, disulfide bond-mediated oligomerization of Death Receptors 4 and 5 (DR4/5) and EGFR, and upregulation of DR5 [2–4, 29]. Therefore, we examined the effect of the novel DDAs on disulfide bonding of these proteins by immunoblot analysis under non-reducing conditions ([4] and Supplemental Information). When applied to cancer cells at the same concentration, dMtcyDTDO and dFtcyDTDO induced more DR4/5 oligomerization, higher DR5 levels, and greater ERS (Fig. 1D, left panel). Induction of ERS by dFtcyDTDO and dMtcyDTDO, as indicated by GRP78 elevation, and disulfide-mediated oligomerization of DR4 and EGFR, were detected at 625 nM and oligomerization of DR5 was observed at 1.25 μM of either dFtcyDTDO or dMtcyDTDO (Fig. 1D, right panel). Additional immunoblot analyses performed under reducing conditions revealed that dMtcyDTDO is more potent than tcyDTDO in the induction of HER3 downregulation, DR5 upregulation, and ERS as indicated by elevated CHOP and XBP1s (Fig. S1B). Time-course and concentration-response analyses of these endpoints showed that dMtcyDTDO induced these responses as early as four hours after treatment (Fig. S1C).

**Figure 1.**
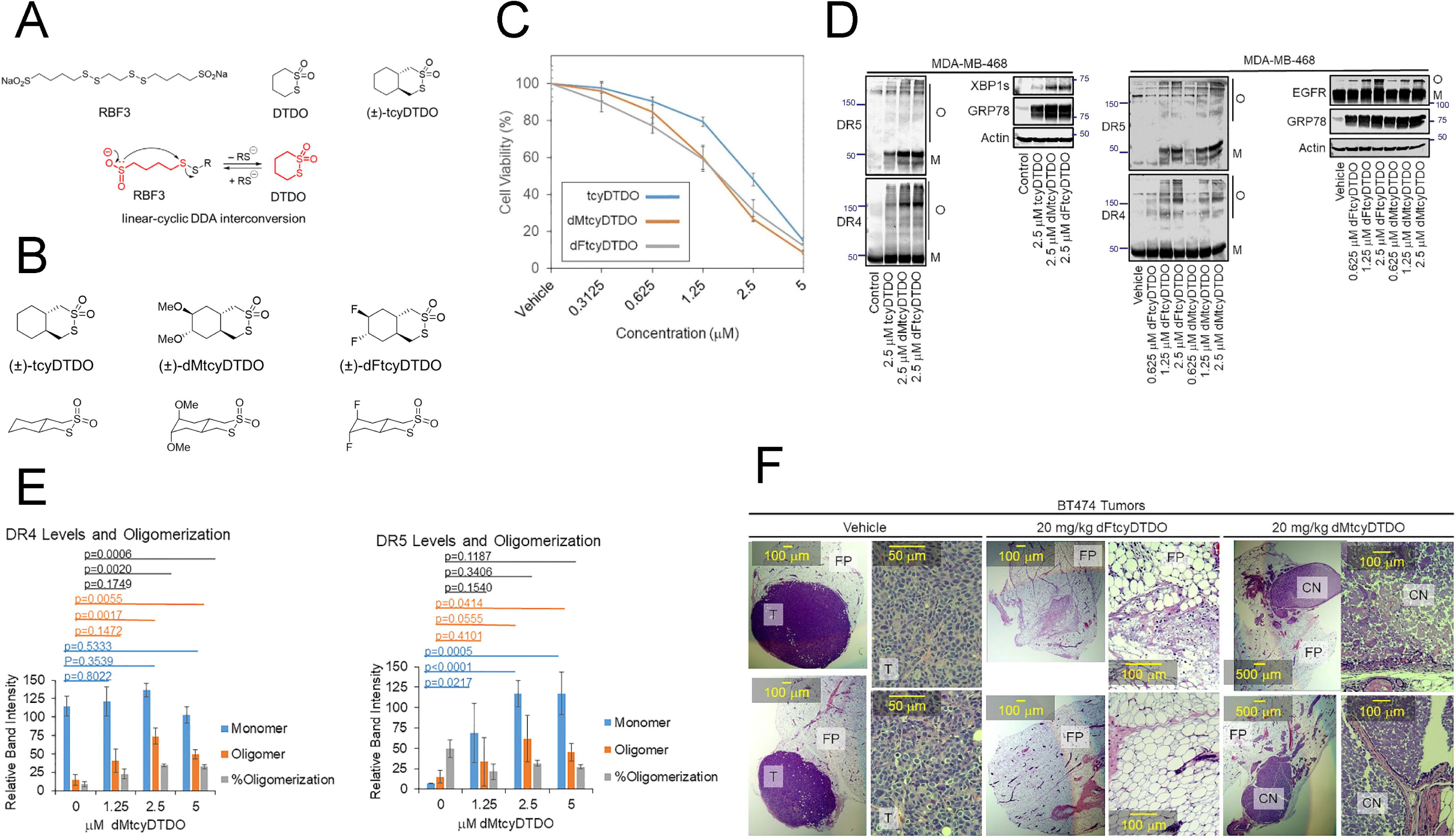
Characterization of new, more potent, orally available DDAs that induce rapid tumor regression. A. Representative first and second generation DDAs (RBF3, DTDO, and tcyDTDO) and illustration of interconversion between the linear and cyclic forms of the DDA pharmacophore. B. Structures of new tcyDTDO derivatives dMtcyDTDO and dFtcyDTDO. C. MDA-MB-468 cells were treated for 24 h with the indicated concentrations of DDAs and cell viability (MTT) assays were performed. Three biological replicates were performed with similar results. D. MDA-MB-468 cells were treated for 24 h as indicated and subjected to immunoblot analysis after non-reducing SDS-PAGE. O and M represent disulfide bonded oligomeric and monomeric protein forms, respectively. (N=4 for each treatment group with similar within-group results). E. Levels and degree of oligomerization of DR4 (left panel) and DR5 (right panel) were determined by densitometry analysis of bands across multiple experiments and subjected to statistical analysis. Values plotted represent averages and error bars represent S.E. P-values for pairwise comparisons were determined using Student’s t-test. F. Mice bearing BT474 cell xenograft tumors were treated once daily for five days with vehicle, 20 mg/kg dFtcyDTDO, or 20 mg/kg dMtcyDTDO. Two hours after the fifth treatment, mammary fat pads/tumors were excised and tumor sections were stained with H&E. T denotes tumor tissue, FP indicates mammary fat pads, and CN represents coagulation necrosis.

Quantitation of dMtcyDTDO effects on the levels and state of oligomerization of DR4 and DR5 revealed that dMtcyDTDO increases DR4 oligomerization without affecting overall DR4 levels, while dMtcyDTDO increases the levels of both oligomeric and monomeric forms of DR5 (Fig. 1E). Tumor studies using the HER2+ BT474 xenograft model [2] were performed to examine if dMtcyDTDO and dFtcyDTDO induce rapid breast cancer cell death in mice bearing palpable breast tumors. Tumors from animals treated with the vehicle for four consecutive days did not show evidence of tumor necrosis (Fig. 1F). In contrast, tumors regressed in mammary fat pads from dFtcyDTDO treated animals. Mammary glands from dMtcyDTDO treated animals exhibited tumors that had undergone coagulation necrosis, consistent with rapid and uniform death of the cancer cells.

### AGR2 as a DDA binding protein

To understand how DDAs kill breast tumors, identification of their direct protein targets is essential. The development of molecular probes proved crucial in identifying DDA biological targets and linking them to the observed DDA responses. For this purpose, we used biotinylated-DDA based affinity pulldown techniques.

To generate DDA probes, we sought a biologically active cyclic DDA with a functional group that enables installation of a biotin tag. This was achieved by fusing a Boc-protected pyrrolidine ring at the 4,5-positions of the parent DTDO compound to generate (±)-BocPyrDTDO (Fig. 2A), which is biologically active and showed a potency similar to DTDO (Fig. S1D). The synthesis of (±)-BocPyrDTDO was accomplished by cleaving the acetyl groups of dithioacetate (±)-**3** followed by oxidation to furnish the 1,2-dithiane 1,1-dioxide ring (Fig. 2A). Intermediate (±)-**3** can be obtained through a few functional group interconversion steps from diester (±)-**1** which itself could be obtained according to literature procedures. The amino group on the pyrrolidine ring, after Boc group deprotection, enables installation of the linker and biotin tag as described in full detail in the Supplemental Information. Biotinylated DDA analogs, including Biotin-PyrDTDO (BPD) and Biotin-GlyPyrDTDO (BGPD), were generated in this fashion and used as affinity tags to identify DDA bound proteins (Fig. 2A).

**Figure 2.**
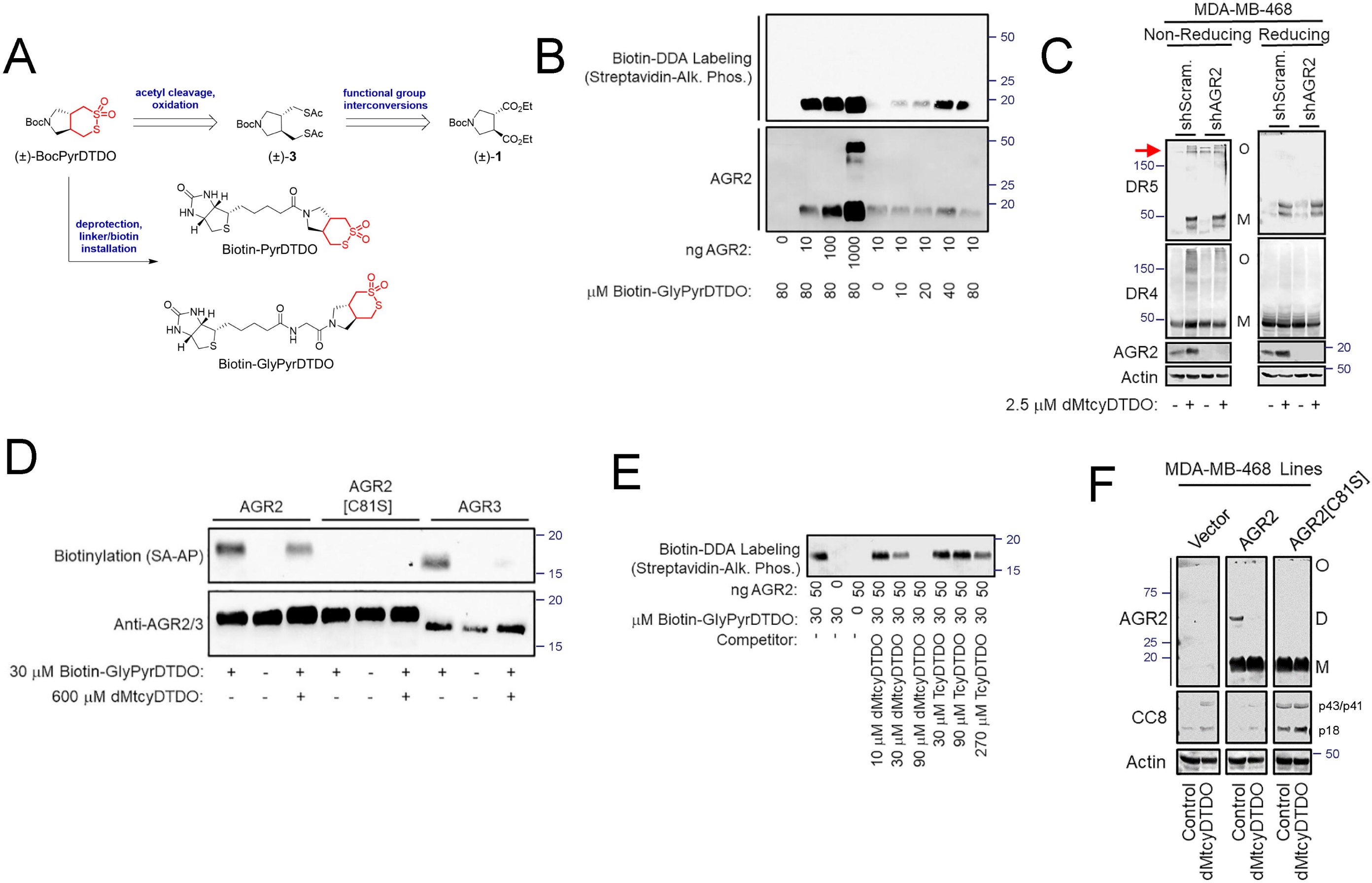
AGR2 and AGR3 are DDA target proteins. A. Abbreviated synthetic approach and chemical structures of the biotinylated DDA analogs Biotin-PyrDTDO and Biotin-GlyPyrDTDO. B. Purified, recombinant AGR2 was reacted with Biotin-GlyPyrDTDO for 1 h at 37°C, resolved by non-reducing SDS-PAGE, transferred to membranes, and probed with Streptavidin-Alkaline Phosphatase, and by anti-AGR2 immunoblot. C. Vector control or AGR2 knockdown MDA-MB-468 cells were treated for 24 h as indicated and analyzed by non-reducing (left panels) or reducing (right panels) immunoblot. Red arrow indicates oligomerized forms of DR5 that are present at higher levels in the AGR2 knockdown cells. O and M represent disulfide-bonded oligomeric and monomeric protein forms, respectively. D. Biotin-GlyPyrDTDO binding assays performed as in Fig. 2B, employing the C81S AGR2 mutant and AGR3 in addition to AGR2. In the indicated samples, the proteins were pre-incubated with 600 μM dMtcyDTDO. E. AGR2 binding assay performed as in Fig. 2B to examine the relative abilities of pre-incubation of AGR2 with dMtcyDTDO or tcyDTDO to block subsequent binding of Biotin-GlyPyrDTDO. F. The indicated MDA-MB-468 stable cell lines were treated for 24 h with 5 μM dMtcyDTDO and subjected to analysis by non-reducing SDS-PAGE/immunoblot. O, D, and M represent the oligomeric, dimeric, and monomeric forms of AGR2 detected. P43/p41 and p18 denote different Caspase 8 cleavage products.

The protein disulfide isomerase (PDI) family member AGR2 was previously implicated in disulfide bonding of EGFR [12], and DDAs alter EGFR disulfide bonding ([4] and Fig. 1D, right panel). We hypothesized that AGR2 might be a target of DDAs and examined if biotinylated DDAs bind covalently with recombinant AGR2. Biotin-GlyPyrDTDO labeled AGR2 in a concentration-dependent manner (Fig. 2B). Under these conditions, Biotin-GlyPyrDTDO labeled monomeric, but not dimeric AGR2. We further hypothesized that in addition to EGFR, DR4 or DR5 may be AGR2 client proteins. Stable knockdown of AGR2 in breast cancer cells increased disulfide-mediated DR5 oligomerization in the absence of DDA treatment (Fig. 2C). AGR2 forms mixed disulfide bonds to its client proteins with Cys81 of its thioredoxin-like repeat [13], and Cys81 is the only Cys residue present in AGR2. Therefore, we examined if mutating Cys81 to Ser prevented DDA labeling. The C81S AGR2 mutant was not labeled by Biotin-GlyPyrDTDO (Fig. 2D). AGR3 is the PDI paralog most closely related to AGR2 and it was also labeled by Biotin-GlyPyrDTDO. Addition of a molar excess of dMtcyDTDO to the labeling reactions partially prevented Biotin-GlyPyrDTDO binding to AGR2 and AGR3 (Fig. 2D), suggesting competition for binding to the same Cys residue(s). A comparison of the ability of dMtcyDTDO and tcyDTDO to block Biotin-GlyPyrDTDO binding to AGR2 indicated that dMtcyDTDO was a more effective competitor, indicating that competition in this binding assay reflects DDA potency in cancer cells (Fig. 2E, S1E, S1F). AGR2 is expressed at low levels in MDA-MB-468 cells, and disulfide-mediated AGR2 complexes with client proteins were not detected in either the presence or absence of dMtcyDTDO treatment (Fig. 2F). Enforced AGR2 expression revealed formation of dimeric AGR2 that was not observed with the C81S AGR2 active site mutant. Treatment of the cells with dMtcyDTDO blocked dimer formation by wild type AGR2, consistent with blockade of C81S by dMtcyDTDO binding.

### Identification of PDIA1 and ERp44 as DDA target proteins

A panel of three breast cancer cell lines was treated with Biotin-PyrDTDO to label DDA target proteins in intact cells. In addition to affinity purification of endogenous biotinylated proteins, such as carboxylases, which use biotin as a cofactor, two biotinylated proteins were observed exclusively in the context of cell labeling with Biotin-PyrDTDO in multiple experiments and termed DDA target proteins 1 and 2 (DDAT1/2). DDAT1/2 migrated at 60 kDa and 44 kDa, respectively, and were observed in three different breast cancer lines in crude cell extracts and in samples purified using Streptavidin-agarose (Fig. 3A). The Streptavidin-agarose purification was scaled up to generate DDAT1/2 in sufficient amounts for identification by mass spectrometry, and aliquots of this preparation and the corresponding crude extracts were analyzed by blotting with Streptavidin-Alkaline Phosphatase, and by silver staining (Fig. 3B). DDAT1 and DDAT2 from a Coomassie stained gel were excised and analyzed by trypsin digestion followed by mass spectrometry as described in Materials and Methods. DDAT1 and DDAT2 were identified as the PDI-family members PDIA1 and ERp44, respectively (Fig. S2A, S2B). In support of these assignments, ERp44, PDIA1, and AGR2 were isolated from the luminal A T47D and luminal B BT474 breast cancer cell lines treated with Biotin-PyrDTDO using Streptavidin-Agarose (Fig. 3C, left panel). In additional Biotin-PyrDTDO pulldown experiments, ERp44 and PDIA1 were detected, but the PDIs ERp57 and ERp5 were not (Fig. 3D). Endogenous proteins are biotinylated via amide linkages. Thus, DDAT1 and DDAT2 were distinguished from endogenous biotinylated proteins by disruption of Biotin-DDA/target protein linkages with the reducing agent 2-Mercaptoethanol (Fig. S2C). Immunoprecipitation with an AGR2 antibody showed that ERp44 and PDIA1 co-purify with AGR2 from cells irrespective of treatment with Biotin-PyrDTDO (Fig. 3C, left panel). ERp44 immunoprecipitates contained PDIA1, but AGR2 co-purification was not detected. Biotin-PyrDTDO/Streptavidin-Agarose also affinity purified ERp44 and PDIA1 from the MDA-MB-468 Triple-Negative breast cancer cell line (Fig. 3C, right panel). In vitro binding studies showed that Biotin-PyrDTDO labeled ERp44 and PDIA1, and in both cases, biotinylation was blocked by pre-incubation with a molar excess of dMtcyDTDO (Fig. 3E). If ERp44 and PDIA1 are cellular DDA targets, then DDA treatment may alter the pattern of ERp44 and PDIA1 disulfide bonding with their client proteins. Consistent with this, treatment of breast cancer cells with dMtcyDTDO reduced disulfide bonding of ERp44 and PDIA1 with multiple proteins (Fig. 3F, red arrows) with minimal effect on monomeric levels of ERp44 and PDIA1. PDIA1 is a validated target for anticancer drugs [16]. However, ERp44 has not been investigated extensively as a therapeutic target for cancer, and small molecule ERp44 inhibitors have not been reported. Small molecule active site AGR2 inhibitors have not been reported, but under the conditions used here, AGR2 binding to client proteins was not detected. Therefore we focused most of our efforts on examining the consequences of ERp44 inhibition by DDAs.

**Figure 3.**
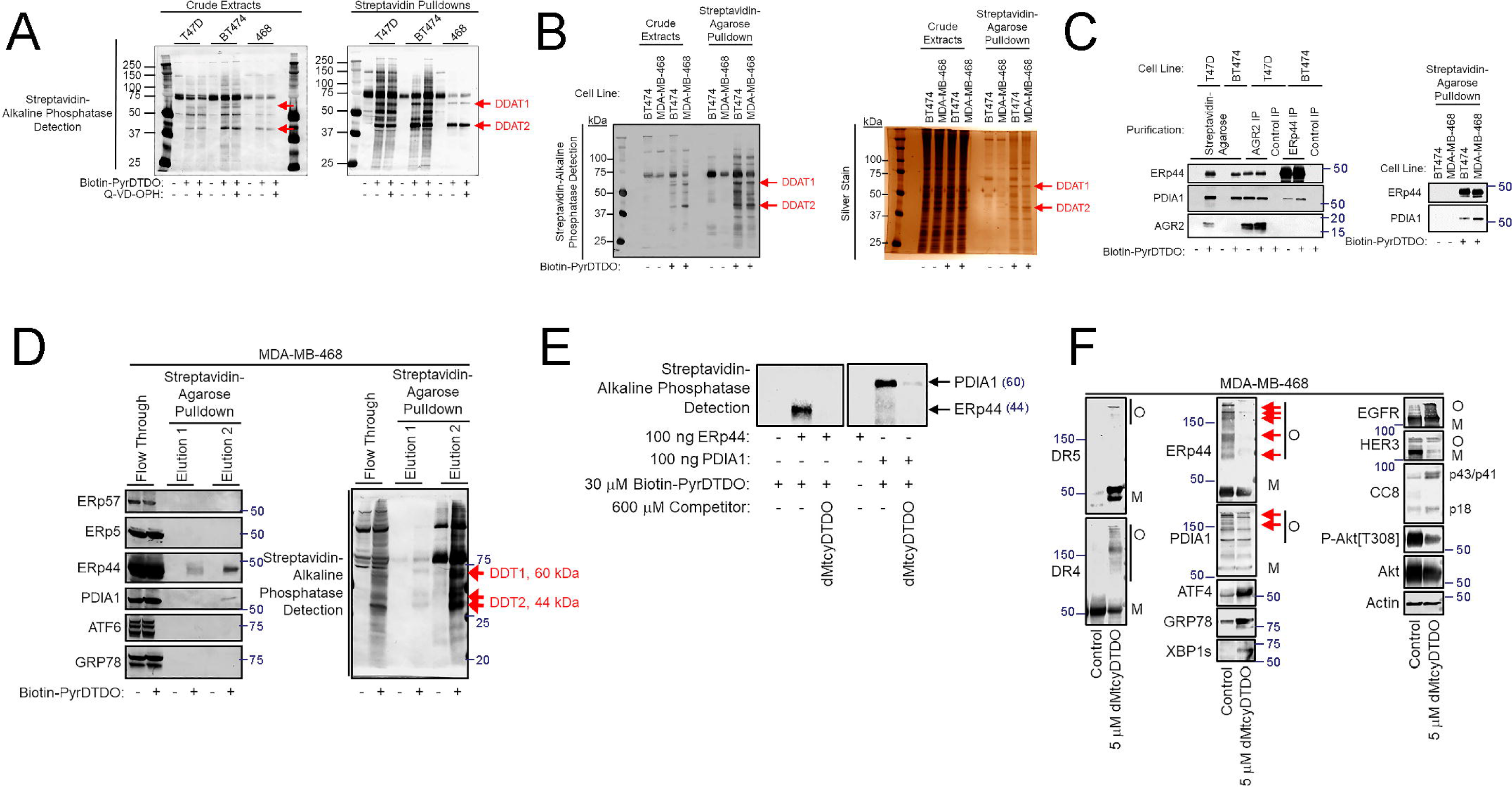
ERp44 and PDIA1 are additional DDA targets. A. Breast cancer cell lines were treated for 16 h with vehicle, 100 μM Biotin-PyrDTDO, or 100 μM Biotin-PyrDTDO + 10 μM Q-VD-OPH for 16 h. Biotinylated proteins were isolated using Streptavidin-coated beads and the crude extracts and Streptavidin pulldowns were analyzed by blotting with Streptavidin-Alkaline Phosphatase. Red arrows indicate bands, designated DDAT1 and DDAT2, present exclusively after Biotin-PyrDTDO treatment. B. Biotin-PyrDTDO/Streptavidin pulldowns were performed in BT474 and MDA-MB-468 cells. Purified material was analyzed by Streptavidin-Alkaline Phosphatase detection and silver stain. C, left panel. The indicated breast cancer lines were treated with vehicle or 100 μM Biotin-PyrDTDO for 16 h and extracts were subjected to Streptavidin-agarose pulldowns, or anti-AGR2 or anti-ERp44 immunoprecipitations. The purified material was analyzed by immunoblot with the indicated antibodies. C, right panel. BT474 and MDA-MB-468 breast cancer lines were treated with vehicle or 100 μM Biotin-PyrDTDO for 16 h and extracts were subjected to Streptavidin-agarose pulldowns. The affinity-purified samples were analyzed by immunoblot. D. MDA-MB-468 cells were treated for 24 h with Biotin-PyrDTDO. Cell extracts were subjected to Streptavidin-agarose pulldowns and the beads were washed and eluted by incubation with 1 M 2-mercaptoethanol (Elution 1) followed by elution with SDS-sample buffer containing 5 mM biotin (Elution 2). The eluents and the flow-throughs were analyzed by immunoblot (left panel) or probed with Streptavidin-conjugated with Alkaline Phosphatase (right panel). E. Binding assays employing recombinant, purified ERp44 and PDIA1. Proteins were labeled with Biotin-PyrDTDO after pretreatment with vehicle or dMtcyDTDO. Reactions were analyzed by non-reducing SDS-PAGE, followed by detection with Streptavidin-Alkaline Phosphatase. F. MDA-MB-468 cells were treated for 24 h with vehicle (control) or 5 μM dMtcyDTDO. Cell extracts were resolved by non-reducing SDS-PAGE and analyzed by immunoblot with the indicated antibodies. O and M represent disulfide bonded oligomeric and monomeric protein forms, respectively (CC8, Cleaved Caspase 8). Red arrows denote bands stained with ERp44 or PDIA1 antibodies that change in intensity with dMtcyDTDO treatment.

### DDAs block the ER retention function of ERp44

ERp44 carries out two important roles by disulfide bonding with client proteins and cycling between the Golgi and the ER as modeled in Fig. 4A. First, ERp44 retrotranslocates inappropriately oligomerized proteins from the Golgi to ER for additional rounds of disulfide bond formation and shuffling. Second, ERp44 forms disulfide bonds with select partner proteins to retain them in the ER/Golgi, preventing their secretion. To determine if DDAs block ERp44 retention of partners by binding to the ERp44 active site, MDA-MB-468 cells were treated with dMtcyDTDO for 24 h and the levels of the ERp44 partner protein ERAP1 were examined. DMtcyDTDO treatment decreased cellular ERAP1 levels (Fig. 4B). Quantitation of dMtcyDTDO effects on ERAP1 levels and dMtcyDTDO binding to its client proteins across multiple experiments showed that dMtcyDTDO decreased ERAP1 levels and ERp44 binding to client proteins in a statistically significant manner (Fig. 4C, D). ERAP1 migrates primarily as a monomer even under nonreducing conditions, indicating that ERAP1 disulfide bonding with ERp44 is transient. Enforced expression of wild type ERp44 (CXXS) resulted in the formation of a higher molecular mass band recognized by ERAP1 antibodies (Fig. 4E, red arrow). This band was not observed when the C58S ERp44 active site mutant (SXXS) was expressed, but was observed when the S61C mutant (CXXC) was expressed. The ERAP1-ERp44 disulfide-bonded dimer was not observed when analyses were performed under reducing conditions (not shown) and was reduced if the cells were treated with the DDA dFtcyDTDO for 24h (Fig. 4E).

**Figure 4.**
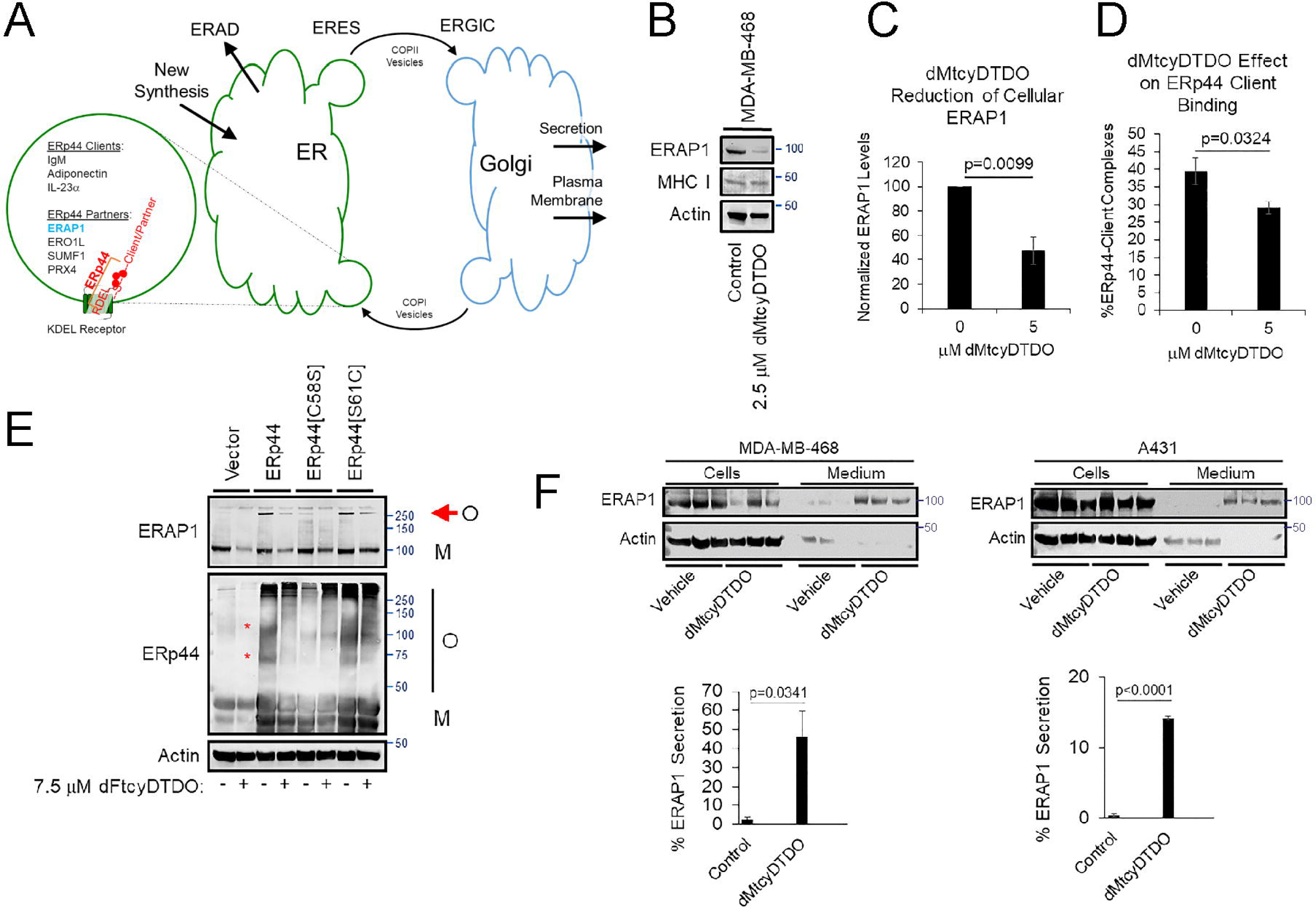
DDAs block ER retention of ERAP1 by ERp44. A. Model for the role of ERp44 in retrotranslocation of client proteins from the Golgi to the ER and retention of partner proteins within the ER. B. Immunoblot analysis of the levels of ERAP1 and MHC I in MDA-MB-468 cells after treatment with dMtcyDTDO for 24 h. C. Quantitative analysis of cellular ERAP1 levels in control and dMtyDTDO treated MDA-MB-468 cells across four experiments. D. Quantitative analysis of cellular ERp44 binding to client proteins in control and dMtyDTDO treated MDA-MB-468 cells across four experiments. E. The indicated stable MDA-MB-468 cell lines were treated for 24 h with dFtcyDTDO and subjected to non-reducing SDS-PAGE followed by immunoblot. O and M represent oligomeric and monomeric protein isoforms. F. MDA-MB-468 (left panels) or A431 cells (right panels) were treated for 24 h with 5 μM dMtcyDTDO and cell extracts and conditioned culture media were subjected to immunoblot analysis as indicated. ERAP1 levels in the cell extracts and culture medium were subjected to statistical analysis (lower panels, N=3).

DDA-induced secretion of ERAP1 was examined by treating MDA-MB-468 breast cancer cells or A431 skin cancer cells with dMtcyDTDO for 24 h and examining ERAP1 levels in cell extracts and in the culture medium. ERAP1 levels in the medium dramatically increased upon dMtcyDTDO treatment (Fig. 4F, upper panels), and dMtcyDTDO induced ERAP1 secretion in a statistically significant manner (Fig. 4F, lower panels).

### A role for ERp44 in the control of Death Receptor and EGFR disulfide bonding

Immunoblot analysis of a panel of breast cancer cell lines revealed that PDIA1 levels were similar among the lines (Fig. 5A). In contrast, ERp44, AGR2, and AGR3 levels varied among the lines, as did the oxidoreductase ERO1, which plays a key role in producing oxidizing equivalents for disulfide bond formation [16, 34–36]. A BT474 breast cancer cell line stably overexpressing ERp44 was generated to examine the effect of elevated ERp44 levels on DR4/5 disulfide bonding and DDA responses. ERp44 overexpression reduced DDA induction of high molecular mass DR4 oligomers, and to a lesser extent high molecular mass DR5 oligomers (Fig. 5B, left panel, red arrows). In both cases, the overall pattern was a shift from high to lower molecular mass disulfide-bonded species. These oligomeric protein forms were not observed when the analyses were performed under reducing conditions (Fig. 5B, right panel). Next, ERp44 knockdown MDA-MB-468 cells were generated to examine the effect of ERp44 depletion on DR4/5 disulfide bonding. Although we were only able to achieve modest ERp44 knockdown, in multiple independent experiments ERp44 knockdown consistently increased high molecular mass oligomers of DR4, DR5, and EGFR in the absence of DDA treatment (Fig. 5C, red arrows; expanded view in Fig. 5D, red asterisks; Fig. S2D). ERp44 knockdown augmented PDIA1 oligomerization and increased dMtcyDTDO induction of Caspase 8 cleavage/activation (CC8). Together, the results of the AGR2 and ERp44 knockdown experiments are consistent with roles for AGR2 and ERp44 in facilitating native disulfide bond formation of DR4 and DR5, and for DDAs in blocking these AGR2 and ERp44 functions. The observation that ERp44 knockdown increases PDIA1 binding to clients (Fig. 5C, 5D) and that ERp44 and PDIA1 form complexes (Fig. 3C) may indicate that PDIA1 and ERp44 chaperone native disulfide bond formation in a coordinated manner.

**Figure 5.**
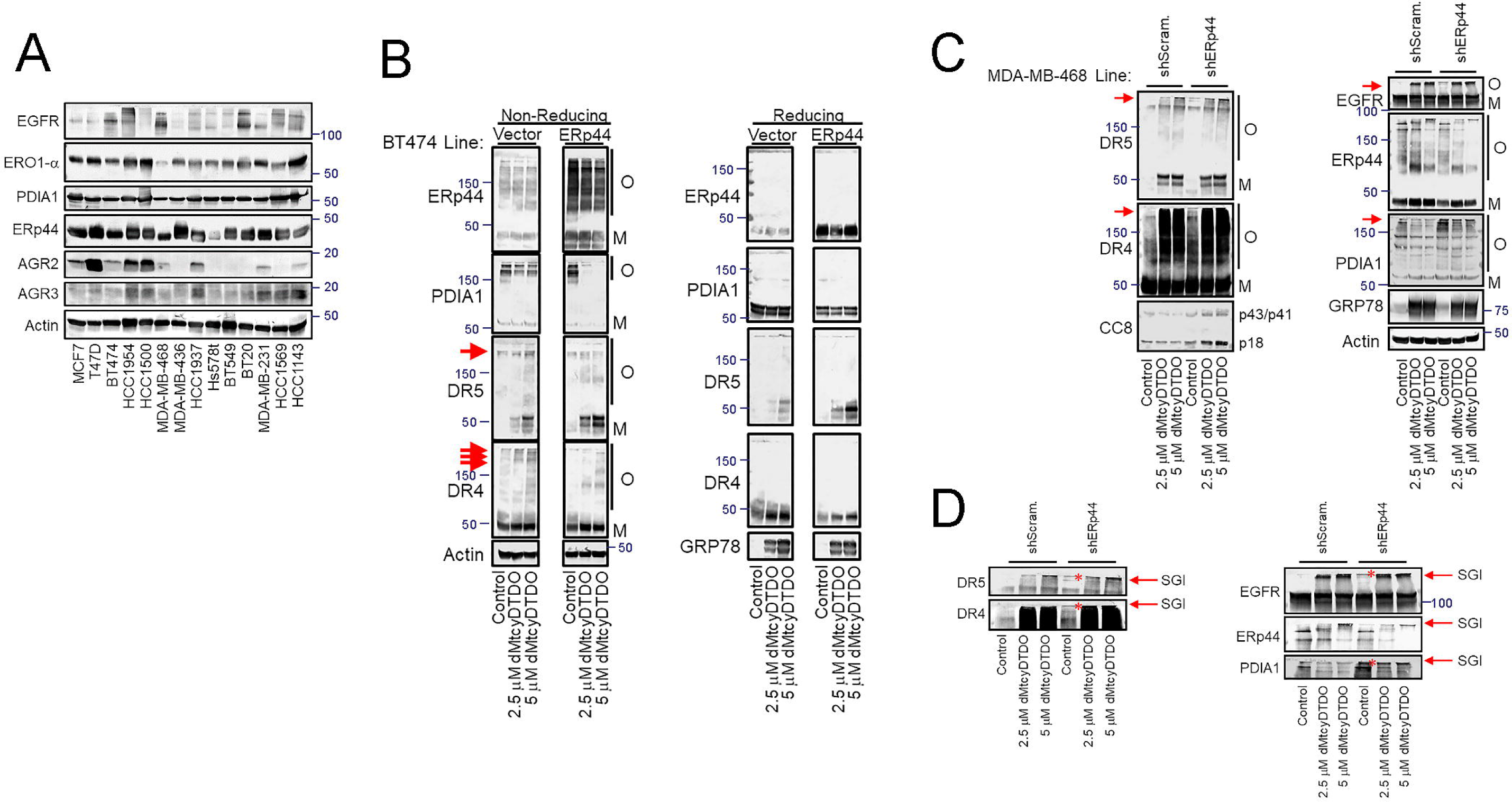
Evidence that disulfide bonding of DR4 and DR5 are regulated by ERp44. A. Immunoblot analysis of the expression levels of EGFR, ERO1-α, AGR2, AGR3, PDIA1, and ERp44 across a panel of breast cancer cell lines. Actin serves as a loading control. B. BT474 cells stably expressing ERp44 or transduced with the vector control were treated for 24 h with vehicle, 2.5 μM, or 5 μM dMtcyDTDO. Cell extracts were resolved by SDS-PAGE under non-reducing (left panel) or reducing (right panel) conditions and analyzed by immunoblot. O and M are disulfide bonded oligomeric and monomeric protein forms, respectively. Red arrows highlight dMtcyDTDO-induced changes in DR4 and DR5 oligomerization. C. Vector control or ERp44 knockdown MDA-MB-468 cell lines were treated for 24 h with vehicle, 2.5 μM, or 5 μM dMtcyDTDO. Cell extracts were separated by non-reducing SDS-PAGE and analyzed by immunoblot (CC8, Cleaved Caspase 8). O and M are disulfide bonded oligomeric and monomeric protein bands, respectively. D. Expanded portions of the blots from panel C showing high molecular mass bands at the stacking gel/resolving gel interface (SGI). Asterisks denote bands at the SGI that are more prominent in the ERp44 knockdown cells in the absence of dMtcyDTDO treatment.

We next examined if mutations within the thioredoxin-like repeat of ERp44 alter its disulfide bonding in the presence or absence of DDA treatment. Overexpressed, wild type ERp44 exhibited a high level of disulfide bond-mediated oligomer formation with client proteins (Fig. 6A). Both endogenous and overexpressed ERp44 monomers migrated as a doublet, which may result from differential O-linked glycosylation [37]. The C58S ERp44 mutant (SXXS), which lacks a Cys residue in its thioredoxin-like repeat, showed lower binding to client proteins at similar expression levels. The S61C ERp44 mutant (CXXC) exhibited slightly increased oligomerization compared with wild type ERp44.

**Figure 6.**
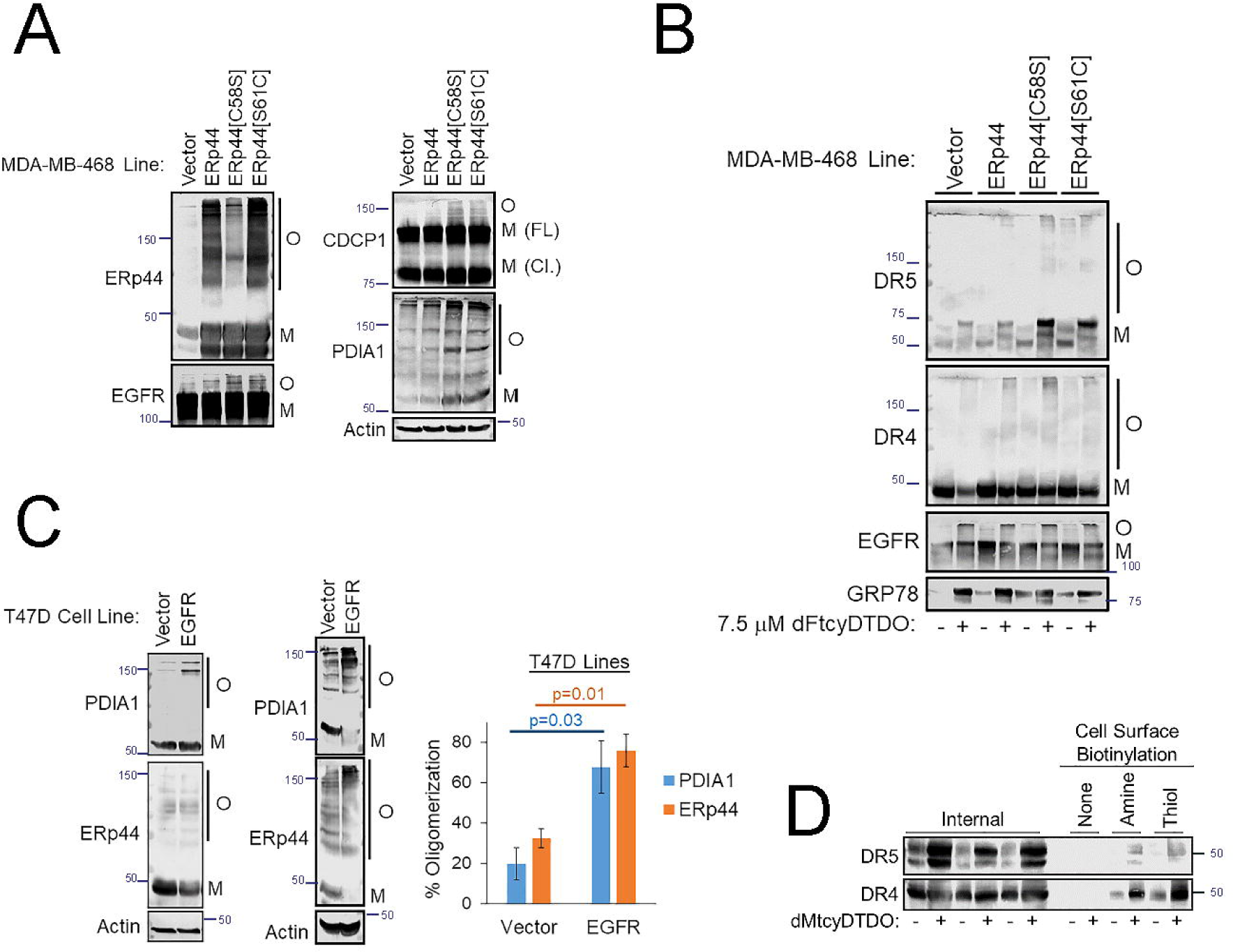
Expression of ERp44 thioredoxin-like repeat mutants alters DDA responses. A. Non-reducing immunoblot analysis of vector control MDA-MB-468 cells, or cells overexpressing wild type ERp44, ERp44[C58S], or ERp44[S61C]. O and M are disulfide bonded oligomeric and monomeric protein bands, respectively. M (FL) and M (Cl.) represent the monomeric full-length and cleaved forms of CDCP1, respectively. B. Non-reducing immunoblot analysis of the samples from Fig. 4E with the indicated antibodies. O and M are disulfide bonded oligomeric and monomeric protein bands, respectively. C, left panel. Extracts from vector control or EGFR overexpressing T47D cells were analyzed by non-reducing SDS-PAGE followed by immunoblot. O and M are oligomeric and monomeric protein bands, respectively. Two replicate experiments are shown involving independent isolation of the T47D vector control and EGFR overexpressing lines. C, right panel. Quantitation of the percentage of PDIA1 and ERp44 present in oligomeric or monomeric forms as assessed by immunoblot and densitometry analysis. Graphic representation of the results of three experiments using independent isolates of T47D/Vector and T47D/EGFR lines examining levels of monomeric and oligomeric PDIA1 and ERp44. Results are shown as the mean ± standard error. D. MDA-MB-468 cells were treated for 24 h with vehicle or 2.5 μM dMtcyDTDO. The cells were then subjected to cell surface protein labeling using membrane-impermeable amine- or thiol-reactive biotinylation probes as described in Supplemental Information. Surface proteins were collected using Streptavidin-coated beads and analyzed along with the flow-though (internal, non-biotinylated proteins) by immunoblot.

We previously showed that DDAs alter the electrophoretic mobility of EGFR and CUB Domain-Containing Protein 1 (CDCP1), block the formation of complexes between these two proteins, and alter their tyrosine phosphorylation patterns [38]. Overexpression of both ERp44 point mutants was associated with increased oligomerization of EGFR and CDCP1, but did not alter the ratio between the full length (FL) and cleaved (Cl.) forms of CDCP1. Analysis of the samples from Fig. 4E for DR4/5 oligomerization revealed that the C58S ERp44 mutant induced the strongest potentiation of DR5 upregulation by dFtcyDTDO (Fig. 6B). Expression of this mutant was also associated with partial oligomerization of EGFR in vehicle treated cells that correlated with increased expression of the ERS marker GRP78 (Fig. 6B). Overall, these observations indicate that ERp44^C58S^, and to a lesser extent, ERp44^S61C^, interfere with disulfide bond formation of ERp44 clients, resulting in higher baseline ERS and potentiating DDA effects on DR4 and DR5. However, these observations do not explain why cancer cells that overexpress EGFR or HER2 have heightened DDA sensitivity [2, 3].

T47D breast cancer cells express low levels of EGFR, HER2, and HER3 and were previously used as a model system to study the effects of EGFR overexpression on DDA responses [2, 3, 39]. Examination of ERp44 and PDIA1 disulfide bonding in control and EGFR overexpressing T47D cells showed that EGFR overexpression resulted in recruitment of ERp44 and PDIA1 into disulfide-bonded protein complexes (Fig. 6C, left panels). Quantitation of the percentages of oligomeric ERp44 and PDIA1 in three experiments employing independently derived pairs of vector control and EGFR overexpressing T47D cell lines showed that EGFR overexpression increased ERp44 and PDIA1 oligomerization (Fig. 6C, right panel). Oligomeric, disulfide-bonded forms of ERp44 and PDIA1 are expected to be inaccessible for binding to newly produced client proteins, while the monomeric forms of ERp44 and PDIA1 are accessible to facilitate native disulfide bonding of their client proteins. Thus, EGFR overexpression may increase DDA sensitivity by reducing the pools of monomeric ERp44 and PDIA1 that DDAs must inhibit in order to block native disulfide bond formation of ERp44 and PDIA1 client proteins.

In addition to chaperoning DR4 and DR5 disulfide bonding, we hypothesized that ERp44, PDIA1, or AGR2 disulfide bonding with DR4 or DR5 may sequester DR4 and DR5 in the ER or Golgi, preventing their trafficking to the cell surface. If so, blocking active site ERp44, PDIA1, and AGR2 Cys residues, DDA treatment may permit DR4 and DR5 trafficking to the plasma membrane. Cell labeling using membrane impermeable, protein-reactive probes is useful for confirming the presence of receptors at the cell surface. Surface biotinylation studies performed on cells after treatment with dMtcyDTDO showed increased cell surface levels of DR4 and DR5, irrespective of whether labeling was performed with amine- or thiol-reactive cell-impermeable biotinylation probes (Fig. 6D). This increased accumulation of DR4/5 at the cell surface may explain previous findings showing that DDAs synergize with the DR4/5 ligand TRAIL to induce cancer cell apoptosis [4].

### DDAs potentiate non-covalent binding of DR5 to ERp44

Data in Fig. 5 and 6 showing that ERp44 knockdown and overexpression of ERp44 mutants alter DR5 levels or oligomerization suggested that DR5 may be an ERp44 client protein. We reasoned that if DR5 is an ERp44 client, the two proteins may bind to each other. ERp44 co-immunoprecipitation experiments showed that ERp44 associates with DR5 and to a lesser extent DR4 in MDA-MB-468 cells, and these interactions were potentiated by DDA treatment (Fig. 7A; Fig. S2E). Unexpectedly, these were non-covalent ERp44/DR5 complexes because although the analyses were performed under non-reducing conditions, each protein migrated as a monomer. ERp44 co-immunoprecipitation studies were next carried out in cells ectopically expressing ERp44 or the C58S or S61C ERp44 mutants. DR5 immunoprecipitated with wild type ERp44 and the S61C ERp44 mutant, but not the C58S ERp44 mutant (Fig. 7B, left panel). DDA treatment increased DR5 and decreased PDIA1 association with ERp44, which may be partly explained by changes in PDIA1 and ERp44 expression levels (Fig. 7B, right panel). Since DDAs increase DR5 levels, which could account for the increased DR5 binding to ERp44, we used the tet-ON system to drive transient high-level DR4 or DR5 expression in the absence of DDA treatment. Doxycycline-induced DR4 expression increased DR4 binding to ERp44 and binding was further enhanced by DDA treatment (Fig. 7C). Doxycycline-induced DR5 expression caused detectable DR5 binding to ERp44, which was strongly enhanced by DDA treatment. The results in Fig. 7A-C demonstrate that DR5 forms a non-covalent complex with ERp44 and that formation of this complex is strongly enhanced by DDA treatment. To determine if ERp44 in dMtcyDTDO-treated cells associates with the disulfide bond-rich extracellular domain of DR5, ERp44 immunoprecipitation experiments were performed in cells that inducibly express wild type DR5 or the Δ81-178 DR5 deletion construct lacking part of the disulfide-rich region. The results indicate that while wild type DR5 bound ERp44 in the context of dMtcyDTDO treatment, ERp44 binding to DR5[Δ81-178] was barely detectable (Fig. 7D). This result suggests that DDA binding to the ERp44 active site Cys residue not only inactivates its catalytic function, but increases its association with the disulfide-rich region of DR5. These catalytically null ERp44(DDA)-DR4/5 complexes may sequester DR4/5, preventing catalytically active forms of ERp44, PDIA1, or AGR2 from chaperoning native DR5 disulfide bond formation.

**Figure 7.**
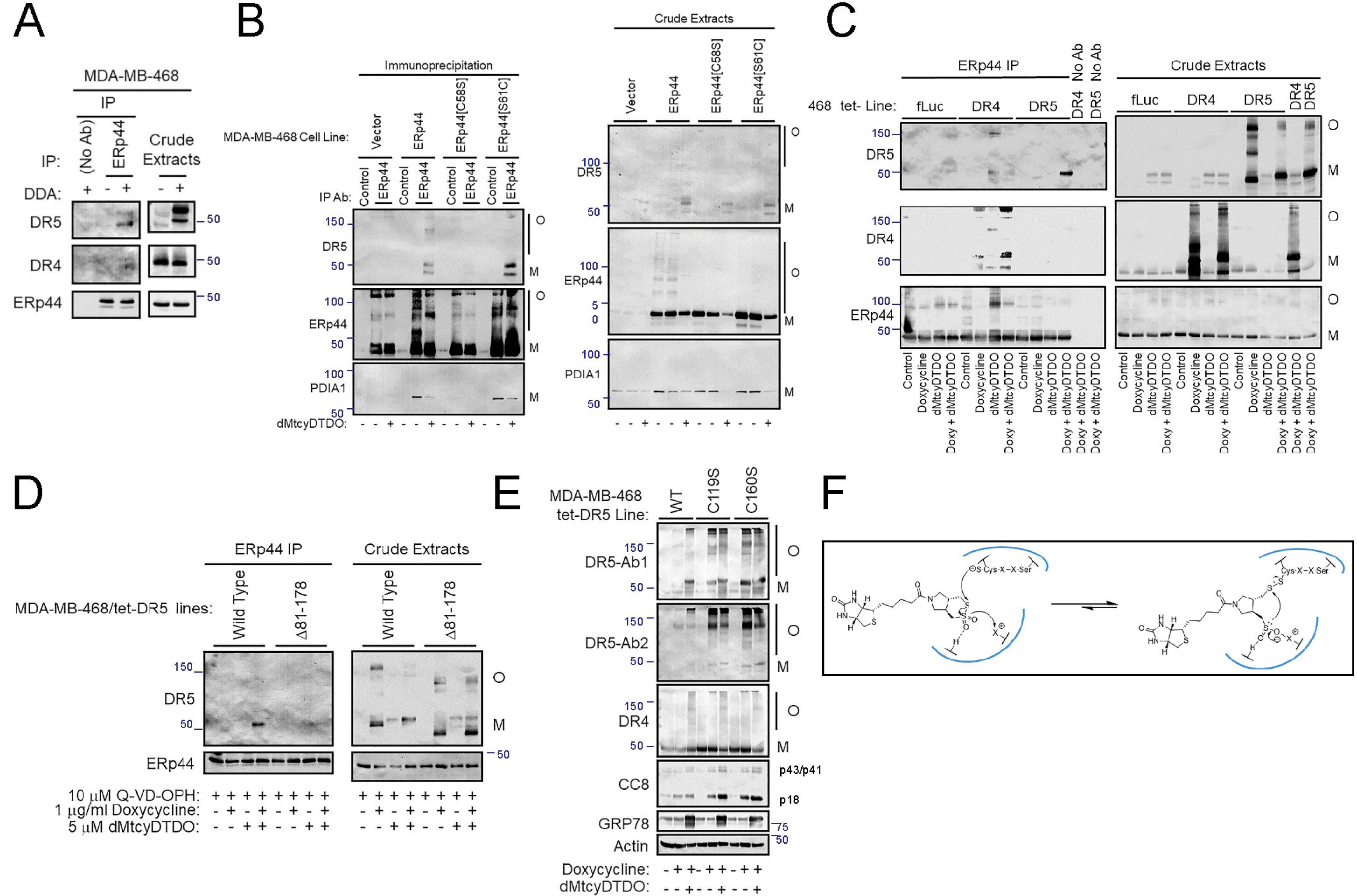
Scission of DR5 disulfide bonds triggers high-level DR5 expression, oligomerization, and Caspase 8 activation. A. MDA-MB-468 cells were treated for 24 h with 10 μM dMtcyDTDO or vehicle and cell extracts were immunoprecipitated with ERp44 antibody and analyzed by immunoblot under non-reducing conditions. B. MDA-MB-468 cells ectopically expressing wild type ERp44 or the C58S or S61C ERp44 mutants were treated with 10 μM dMtcyDTDO or vehicle. Left panels, cell extracts were immunoprecipitated with ERp44 antibody and analyzed by immunoblot. Right panels, crude extracts were analyzed for protein expression levels and oligomerization by immunoblot. O and M represent disulfide bonded oligomeric and monomeric protein forms. C. MDA-MB-468 expressing doxycycline-inducible DR4 and DR5 were treated with 1 μg/ml doxycycline, 10 μM dMtcyDTDO, or doxycycline + dMtcyDTDO for 24 h followed by immunoprecipitation of ERp44 and immunoblot analysis of the immunoprecipitates and crude lysates. O and M represent disulfide bonded oligomeric and monomeric protein forms. D. The indicated MDA-MB-468 cell lines stably expressing wild type DR5 or the Δ81-178 mutant lacking part of the disulfide-rich extracellular domain were treated for 24 h as indicated and subjected to immunoprecipitation with ERp44 antibodies. The immunoprecipitates and crude cell lysates were analyzed by non-reducing SDS-PAGE/immunoblot. O and M represent disulfide bonded oligomeric and monomeric protein forms. E. The indicated stable MDA-MB-468 cell lines were treated for 24 h as indicated and subjected to immunoblot analysis after resolving proteins by non-reducing SDS-PAGE. O and M represent disulfide-bonded oligomeric and monomeric protein forms. F. Model for how the biotin-labelled DDA biotin-PyrDTDO reacts covalently with the PDI active site thiolate. Covalent DDA binding is associated with ring-opening and exposure of a nucleophilic sulfinate group, which is capable of attacking the DDA-PDI disulfide bond, resulting in ring closure and DDA excision. It is hypothesized that solvation of the DDA sulfinate group by the PDI slows the rate of DDA self-excision.

The results presented thus far indicate that DDAs alter the disulfide bonding status of DR4 and DR5, which correlates with inhibition of ERp44 and PDIA1 binding to their client proteins. However, it is unclear if genetic perturbation of DR5 disulfide bonding mimics the effects of DDAs on DR5, which include upregulation of DR5 protein levels and increased pro-apoptotic signaling. This issue was addressed by inducibly expressing DR5 mutants in which a Cys residue was mutated to Ser to disrupt one of the seven extracellular DR5 disulfide bonds. The C119S mutant was constructed to disrupt the center/fourth disulfide bond, and the C160S mutant was constructed to disrupt the seventh, most C-terminal DR5 disulfide bond. As observed previously using this doxycycline-inducible expression system [39], induction of wild type DR5 expression weakly increased DR5 expression (Fig. 7E). However, dMtcyDTDO/doxycycline co-treatment further increased DR5 expression, demonstrating DR5 stabilization by DDA treatment. The C119S and C160S DR5 mutants were highly expressed in both the monomeric and oligomeric forms, and this was associated with Caspase 8 cleavage/activation. DMtcyDTDO treatment modestly increased Caspase 8 cleavage, but did not further increase expression of the C119S or C160S DR5 mutants. Expression of the DR5 mutants did not upregulate the ERS marker GRP78, indicating that the mutation-driven upregulation of DR5 was not a result of ER stress induction. These results indicate that disruption of individual DR5 disulfide bonds is sufficient to promote high-level DR5 protein expression and pro-apoptotic signaling that is not further enhanced by DDA treatment.

## Discussion

### Models for DDA mechanisms of action and compound selectivity

The findings presented here suggest that DDAs bind to the PDI-family proteins ERp44, PDIA1, AGR2, and AGR3 and block their ability to chaperone the disulfide bonding of subsets of client proteins. The resulting alteration of disulfide bonding causes upregulation of DR5, oligomerization of DR4 and DR5, and accumulation of DR4 and DR5 at the cell surface. Oligomerization of DR4 and DR5 causes Caspase 8 activation, which in turn activates Caspase 3 to mediate apoptosis. Caspase activation plays a key role in downregulating EGFR and HER2, and decreasing Akt phosphorylation because these DDA responses are largely ablated by a pan-Caspase inhibitor [4]. To our knowledge, DDAs are the first small molecule agents that block the active site Cys residues of ERp44 and AGR2/3. Further, DDAs are the first experimental anticancer agents that trigger DR4 and DR5-mediated Caspase 8 activation by inducing disulfide bond-mediated DR4/5 clustering in the absence of their ligand, TRAIL. A recent report showed that the extracellular domain of DR5 acts to prevent its oligomerization in the absence of TRAIL [11]. DDAs may overcome this autoinhibition by altering the conformations of the DR4/5 extracellular domains or favoring intermolecular disulfide bonding between DR4/5 extracellular domains over intramolecular disulfide bonding. Consistent with these interpretations, mutational disruption of individual DR5 disulfide bonds is sufficient to trigger DR5 upregulation, oligomerization, and downstream activation of Caspase 8 (Fig. 7E).

Previous reports showed that overexpression of EGFR amplifies multiple DDA responses, including ERS induction, DR5 upregulation, HER1-3 downregulation, and reduction of Akt phosphorylation [2–4]. Discovery of ERp44 and PDIA1 as DDA target proteins led to the observation that EGFR overexpression reduced the fractions of ERp44 and PDIA1 in the monomeric forms and increased the relative abundance of the oligomeric forms. Since ERp44 and PDIA1 oligomerization with client proteins involves their active site Cys residues, only monomeric ERp44 and PDIA1 are accessible for facilitating native disulfide bonding of client proteins. Because DDAs covalently modify monomeric ERp44 and PDIA1 (Fig. 3), these observations may explain the molecular basis for how EGFR overexpression sensitizes cells to DDAs. Given the importance of EGFR as a potential oncogenic driver in a number of cancers, including glioblastoma [40, 41] and Triple-Negative Breast Cancer [42–44], DDAs may have utility against these malignancies.

Because of the relative structural simplicity of DDA molecules, the mechanisms responsible for their target protein selectivity are of interest. PDIA1 has been investigated as a target for anticancer therapeutics, and PDIA1 inhibitors have established anticancer efficacy in preclinical models [45, 46]. AGR2 contributes to tumor progression by facilitating EGFR disulfide bonding and surface presentation [47]. However, AGR2 may also be secreted and promote tumor angiogenesis [48, 49], and act on cell surface receptors [50]. Monoclonal antibodies designed to block extracellular AGR2 functions are under development [50, 51], but agents that ablate the role of AGR2 in the folding of client proteins are not available.

The canonical thioredoxin-like catalytic site sequence CXXC is featured in PDIA1 and several other PDI enzymes (reviewed in [15]). Interestingly, AGR2, AGR3, and ERp44 share a non-canonical CXXS thioredoxin-like repeat sequence. A report employing both yeast genetics and biochemical enzyme assays showed that the essential function of the PDI enzyme for yeast cell survival is catalysis of disulfide exchange or “scramblase” activity and that this could be carried out by the CXXS mutant of PDI (PDI^CXXS^) [17]. Enzyme assays showed that PDI^CXXS^ was able to reactivate inappropriately disulfide bonded RNAse A via its scramblase activity. In contrast, PDI^CXXS^ was unable to catalyze disulfide bond oxidation or reduction like the wild type PDI^CXXC^ enzyme.

The C-terminal Cys residue in the CXXC motif of PDIs accelerates substrate release. Consequently, CXXS substrate-trapping mutants of PDIs have been used to isolate mixed disulfide bonded dimers with their client/substrate proteins [52, 53]. That the CXXS motif is associated with slow client protein off-rate is consistent with the established roles of ERp44 in retaining specific client/partner proteins such as ERO1 [22] and ERAP1 [25] in the ER, and in retrotranslocation of inappropriately disulfide bonded Adiponectin [54, 55] and IgM [21, 56] from the Golgi to the ER (see model in Fig. 4A). Thus, the CXXS motifs of AGR2, AGR3, and ERp44 bound to DDAs may exhibit a slow DDA off-rate for the same reason, and could explain stable DDA binding to these enzymes. Our current working model for DDA binding to AGR2, AGR3, and ERp44 is shown in Fig. 7F. To wit, binding of Biotin-PyrDTDO is initiated by nucleophilic attack of the thiosulfonate group of Biotin-PyrDTDO by the thiolate nucleophile of the Cys residue of the CXXS motif. This results in reversible ring opening. We hypothesize that the rate of ring closure and DDA compound excision is reduced by stabilization of the open form of the DDA by solvation of the sulfinate group through interactions with functional groups in or near the PDI active site. The reason for stable DDA binding to PDIAI, but not ERp57 or ERp5 is unclear since each protein contains two canonical CGHC repeats. Further work is required to determine which PDIA1 Cys residue binds DDAs and to identify features in the local environment that stabilize DDA binding.

It is clear from the experiments with intact cancer cells that DDAs alter the disulfide bonding of DR4, DR5, and EGFR and change the pattern of disulfide bonding between ERp44 and PDIA1 and their client proteins. It is difficult to compare these DDA actions with those of published PDI inhibitors since most of these studies did not examine perturbation of protein disulfide bonding in cancer cells, but rather examined compound activity in the Insulin reduction/aggregation assay. While the Insulin reduction assay is valuable for optimizing the potency of PDI inhibitors, it may not be salient in instances where disulfide isomerase activity, ER retention, or Golgi-ER retrotranslocation are more important than disulfide oxidase or reductase activity.

The presence of 21 different PDIs in humans and the diverse phenotypes of the corresponding knockout mice (e.g., [25, 57–59]) suggests that individual PDIs chaperone the disulfide bonding, ER retention, or Golgi-ER retrotranslocation of different subsets of client proteins. For instance, ERp44 is unique in its ability to retain the aminopeptidase ERAP1 in the ER/Golgi and this results in ERAP1 secretion in ERp44 knockout mice or ERp44-deficient cell lines [25]. A biological consequence of this is that ERp44 deletion is associated with hypotension in mice since secreted ERAP1 destabilizes Angiotensin II, leading to reduced blood pressure. It will be of interest to determine if DDAs exhibit antihypertensive activity.

ERAP1 functions as a “molecular ruler” to process peptides for loading onto MHC I for presentation to T cells [60–62]. ERAP1 inhibitors or ERAP1 knockdown changes the spectrum of the MHC I-associated peptides that constitute the “immunopeptidome.” Since DDAs cause ERAP1 secretion, it will be important to determine if DDAs alter the immunopeptidome, and if so, how this impacts anticancer immunity.

In summary, the results presented here establish perturbation of disulfide bonding by inhibition of subsets of PDIs as a new strategy for activating DR5 and DR4 in cancer cells independently of their engagement by agonistic antibodies or TRAIL. Further, DDAs upregulate DR5 and increase DR4 and DR5 localization to the cell surface. Consequently, DDAs may overcome some of the shortcomings associated with TRAIL analogs and agonistic antibodies specific for DR4 and DR5. Additional research is needed to elucidate the mechanisms by which disrupting individual DR5 disulfide bonds increases its expression levels, disulfide-mediated oligomerization, and activation of Caspase 8. In future studies, it will be important to determine if C-S point mutants of DR4 mimic the DR5 mutants and drive pro-apoptotic signaling. It will also be interesting to examine whether activated point mutants of DR4 and DR5 exhibit the same cancer-selective killing as their ligand, TRAIL.

## Materials and Methods

### In vivo tumor studies and histochemical analysis

Breast tumor formation was initiated in NOD-SCID-gamma (NSG) mice obtained from Jackson Laboratories (Bar Harbor, ME USA) by injecting 1 × 10^6^ cancer cells into the #4 mammary L L fat pads as described previously [4]. Animals bearing HER2+ BT474 tumors [2] were treated with the vehicle or 20 mg/kg dMtcyDTDO or dFtcyDTDO by intraperitoneal injection for four consecutive days. Tissue samples fixed in 4% paraformaldehyde/Phosphate-Buffered Saline (PBS) and paraffin-embedded were sectioned and stained with hematoxylin and eosin (H&E) by the University of Florida Molecular Pathology Core (https://molecular.pathology.ufl.edu/).

### Streptavidin-Agarose Pulldowns

Cells were treated with Biotin-PyrDTDO with or without the pan-Caspase inhibitor Q-VD-OPH to prevent apoptosis (MedChemExpress, Monmouth Junction, NJ USA) for 16 h in 10% FBS– DMEM, followed by the addition of 25 mM *N*-ethylmaleimide (NEM) (Thermo Scientific, Rockford, IL USA) for 20 min at 37°C. Cells were subsequently scraped directly into the media, washed twice with ice-cold PBS, and pelleted by centrifugation. Cells were extracted in buffer (20 mM HEPES (pH 7.4), 1 mM EDTA, 1mM EGTA, 5% Glycerol, 1 nM Microcystin, 1 mM Na_3_VO_4_, and 40 mM Na_2_H_2_P_2_O_7_) containing 1% Triton X-100. Samples were sonicated, followed by centrifugation for 20 min at 4°C. The supernatant was pre-cleared with Sepharose CL-6B (Amersham Biosciences AP, Uppsala, Sweden) for 2h at 4°C. Pre-cleared samples were then incubated with Streptavidin (SA)-Agarose (Thermo Scientific, Rockford IL USA) and mixed overnight at 4°C. The SA-Agarose was washed three times with extraction buffer containing 0.1% Triton X-100, and eluted by boiling in 2X-SDS Sample Buffer containing 5 mM biotin. Samples were analyzed by immunoblot, silver stain, or probing proteins transferred to nitrocellulose membranes with Streptavidin-conjugated Alkaline Phosphatase (Rockland Immunochemicals, Inc., Limerick, PA USA).

### Disulfide Bond-mediated Oligomerization

To evaluate protein disulfide bond-mediated oligomerization, cells were treated with DDAs for 24 h in 10% FBS–DMEM. Cells were subsequently scraped into the media and pelleted by centrifugation. Cell pellets were extracted directly into boiling 2X-SDS-Sample Buffer containing 100 mM NEM and boiled for 10 min. Samples were sonicated, followed by centrifugation for 20 minutes at room temperature, and subsequently analyzed by SDS-PAGE/immunoblot.

### Surface Biotinylation

Cell surface proteins were labeled by incubating cells with 1.6 mM Sulfo-NHS-SS-Biotin (Thermo Fisher Scientific, Waltham, MA USA) or 1.6 mM Biotin-dPEG_3_-Maleimide (MilliporeSigma, St. Louis, MO USA) in PBS, pH 8.0 for 30 minutes at 4 °C. The cells were subsequently washed three times with PBS, and proteins were extracted in 1% Triton X-100 extraction buffer in the absence of reducing agents. Cell extracts were then mixed with Streptavidin-agarose beads (Thermo Fisher Scientific, Waltham, MA USA) for 2.5 h at 4°C. Following centrifugation, the supernatant containing non-biotinylated proteins (flow-through) was retained and boiled in 2X-SDS-Sample Buffer containing 1% 2-mercaptoethanol. Streptavidin-agarose beads bound to biotinylated proteins were washed four times with 0.1% Triton X-100 extraction buffer, and eluted by boiling in 2X-SDS-Sample Buffer containing 1% 2-mercaptoethanol. Samples were then analyzed by immunoblot or blotting with Streptavidin-conjugated Alkaline Phosphatase.

### His_6_-AGR2 TALON Purification

Competent BL21(DE3) cells were transformed with either the AGR2/pET-45b(+) or AGR2 C81S/pET-45b(+) vectors. Protein expression was induced by the addition of 50 μM Isopropyl β-D-1-thiogalactopyranoside (Fisher Scientific, Fair Lawn, NJ USA) to bacterial cell cultures for 2.5 hours with shaking at 37°C. Bacterial cells were subsequently pelleted and dissolved in Mouse Tonicity Phosphate Buffered Saline (MTPBS) containing 1% Triton X-100 and sonicated for 20 seconds on ice, followed by centrifugation for 10 min at 4°C to pellet cell debris. The supernatant was transferred to a new tube, followed by the addition of TALON Resin (Clontech, Mountain View, CA USA) and mixed for 30 min at 4°C. The TALON Resin was pelleted by centrifugation and washed twice with MTPBS and twice with MTPBS containing 30 mM Imidazole. Protein bound to the TALON Resin was eluted with 400 mM Imidazole and precipitated with 60% saturated ammonium sulfate. The precipitated protein was pelleted by centrifugation, dissolved in MTPBS containing 1 mM DTT and 1 mM EDTA, and subsequently dialyzed against a solution of MTPBS containing 10 μM DTT and 30% Glycerol.

### DDA-AGR2 Binding Experiments

Purified His_6_-AGR2 or His_6_-AGR2[C81S], as prepared above, or His_6_-AGR2 (PRO-580) or His_6_-AGR3 (PRO-242) obtained from ProSpec (New Brunswick NJ USA) were incubated with or without varying concentrations of the unlabeled DDA competitors, tcyDTDO or dMtcyDTDO, and 30 μM Biotin-GlyPyrDTDO or Biotin-PyrDTDO in buffer (20 mM HEPES (pH 7.4), 1 mM EDTA, 1mM EGTA, 5% Glycerol, 1 nM Microcystin, 1 mM Na_3_VO_4_, and 40 mM Na_2_H_2_P_2_O_7_) containing 1% Triton-X 100 for 30 min at 37°C. Preliminary studies showed that Biotin-GlyPyrDTDO or Biotin-PyrDTDO exhibited indistinguishable labeling of proteins in intact cells and purified proteins in vitro, and are referred to collectively as “Biotin-DDA” and the two probes were used interchangeably. Reactions were stopped by boiling the samples in 2X-SDS-Sample Buffer containing 100 mM NEM for 10 min. Biotin-DDA labeling of protein was verified by blotting with Streptavidin-Alkaline Phosphatase. ERp44 (PRO-547) and PDIA1 (ENZ-262) were obtained from ProSpec and used in experiments examining the binding of these proteins to Biotin-PyrDTDO.

### Quantitative Analysis of Immunoblot Results

Immunoblots were scanned, converted to grayscale and inverted using Adobe Photoshop (Berkeley, CA). Bands were quantified using ImageJ (NIH, Bethesda, MD). Expression levels were normalized using Actin as a loading control.

### Chemical Synthesis of DDAs

Full experimental and characterization details associated with the synthesis of (±)-BocPyrDTDO, Biotin-GlyPyrDTDO, and Biotin-PyrDTDO are provided in the Supplemental Information. Detailed discussion of the syntheses and analytical characterization of dMtcyDTDO and dFtcyDTDO will be published elsewhere.

### Detailed Methods in Supplemental Information

At least three biological replicates of each of the studies presented were performed. Please see Supplemental Information for in-depth descriptions of approaches relating to cell culture, preparation of cell extracts, immunoblot analysis, construction of retroviral and lentiviral vectors and their use to generate stable cell lines, DNA and protein synthesis assays, cell viability assays, sample preparation for analysis by tandem mass spectrometry, and statistical methods..

## Acknowledgements

These studies were supported in part by grants from the Florida Breast Cancer Foundation (BL and RC), the Ocala Royal Dames for Cancer Research (BL), and the Office of the Assistant Secretary of Defense for Health Affairs through the Breast Cancer Research Program under Award Nos. W81XWH-15-1-0199 (BL) and W81XWH-15-1-0200 (RC), and NIH/NCI grant CA252400 (BL). RF is grateful to the University of Florida for a Graduate School Fellowship. EY thanks the Dr. Howard and Brenda Sheridan Fund in Chemistry for a 2020 Howard and Brenda Sheridan Summer Fellowship. AG is grateful to the Tarrant Organic Chemistry Fund for a 2021 Tarrant Summer Graduate Research Fellowship. Mass spectrometric data on DDA protein targets were obtained by the Proteomics and Mass Spectrometry Facility at the Interdisciplinary Center for Biotechnology Research. Mass spectrometric data on DDA compound structures and DDA adducts were obtained by the UF Department of Chemistry Mass Spectrometry Research and Education Center supported, in part, by the National Institutes of Health (NIH S10 OD021758-01A1).

