## Supplemental Information for "Inhibitors of ERp44, PDIA1, and AGR2 induce disulfide-mediated oligomerization of Death Receptors 4 and 5 and cancer cell death"

**Affiliations**
[1]Department of Pharmacology & Therapeutics, University of Florida, Gainesville, FL 32610 USA.
[2]Department of Chemistry, University of Florida, Gainesville, FL 32611 USA.
[3]Proteomics and Mass Spectrometry Facility, Interdisciplinary Center for Biotechnology Research, University of Florida, Gainesville, FL 32610 USA.
[4]Department of Biology, Genetics Institute, University of Florida, Gainesville, FL 32610 USA
[5]Institute of Molecular Medicine, College of Medicine and Center for Infectious Disease and Signaling Research, National Cheng Kung University, Tainan, Taiwan.
[6]Department of Medicine, University of Florida, Gainesville, FL 32610 USA.
[7]UF-Health Cancer Center, University of Florida, Gainesville, FL 32610 USA.
[8]Department of Pathology, Copenhagen University Hospital Herlev, DK-2730 Herlev, Denmark.

[#]Co-Contributing Corresponding Authors

Table of Contents — Pages

Supplemental Data
Supplemental figure (S) 1: Characterization of novel DDAs and an AGR2 DDA binding assay 2
S2: Identification of ERp44 and PDIA1 as DDA targets 3-11

Cell Culture and Mass Spectrometry Analyses
Cell culture and Immunoblot Analysis 12
Construction of Recombinant Retroviral and Lentiviral shRNA Vectors 12-13
Thymidine Incorporation Assays 14
Protein Synthesis Assays 14
Cell Viability Assays 14
Construction of AGR2 Vectors 14-15
Sample Preparation and Analysis by Mass Spectrometry 15-17
Statistics 17
Literature Cited 17-18

Chemical Syntheses
General Methods 19
Synthesis of (±)-BocPyrDTDO 19-21
Synthesis of BioGlyPyrDTDO (BGPD) 21-23
Synthesis of Biotin-PyrDTDO (BPD) 23
Compound Analysis by NMR 24-34
Literature Cited 34

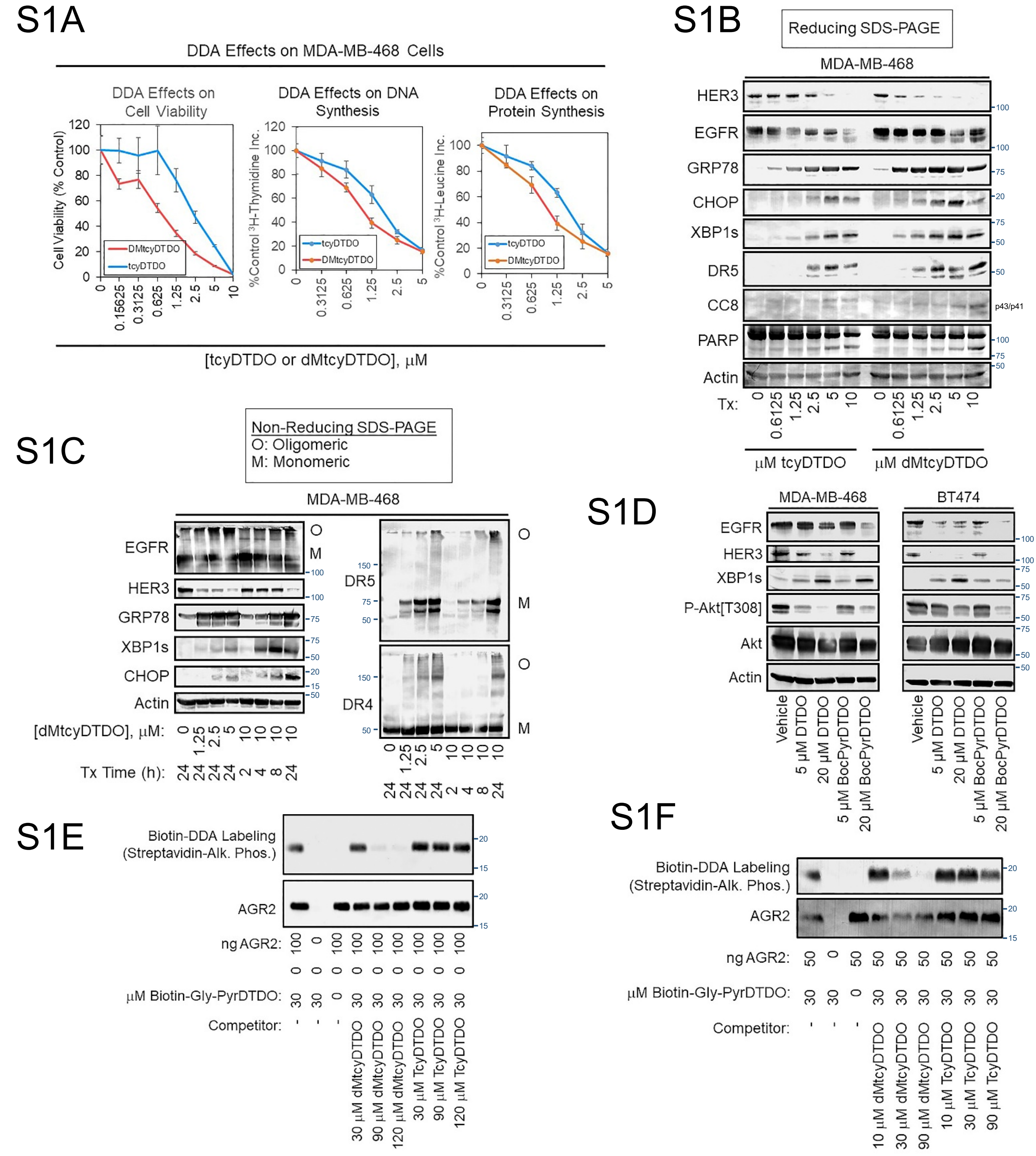


Fig. S1-Characterization of novel DDAs and AGR2 DDA binding assays. S1A. MDA-MB-468 cells were treated for 24 h and assayed for viability, DNA synthesis, and protein synthesis in MTT assays, and tritiated thymidine and leucine incorporation assays, respectively. S1B. Reducing immunoblot analysis of MDA-MB-468 cells treated for 24 h as indicated. S1C. Non-reducing immunoblot analysis of MDA-MB-468 cells treated for 24 h as indicated. S1D. Reducing immunoblot analysis of MDA-MB-468 and BT474 cells treated for 24 h as indicated. S1E. and S1F. AGR2 binding assays employing recombinant AGR2 and Biotin-GlyPyrDTDO under the indicated conditions, followed by immunoblot analysis for AGR2 or probing with Streptavidin-conjugated Alkaline Phosphatase.

# S2A

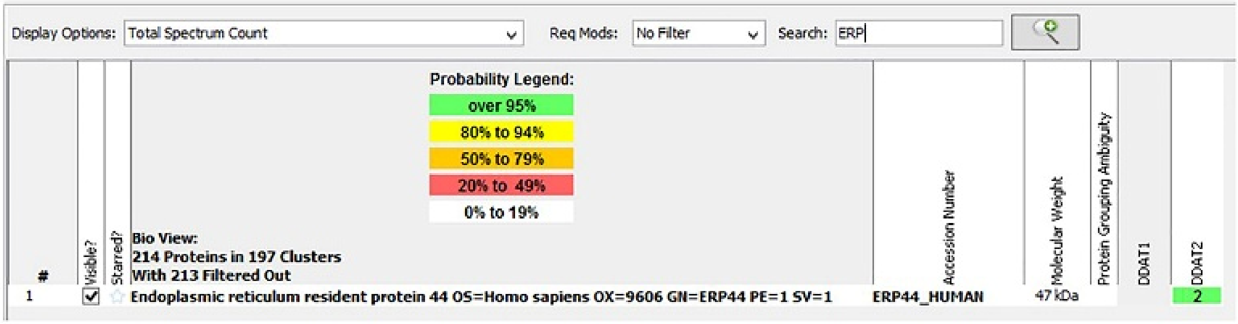


Unique Peptide Search: ERp44 (one peptide sequence; two peptides)

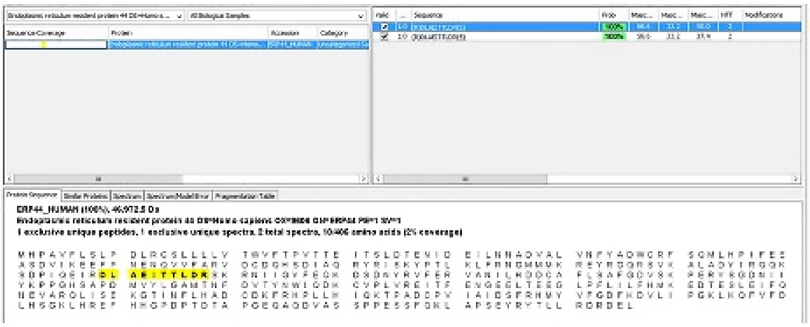

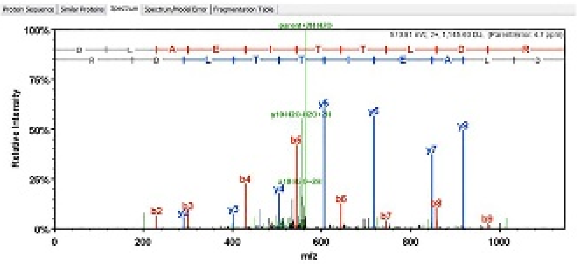

| B | B Ions | B+2H | B-NH3 | B-H2O | AA | Y Ions | Y+2H | Y-NH3 | Y-H2O | Y |
|---|---|---|---|---|---|---|---|---|---|---|
| 1 | 116.0 | | | 98.0 | D | 1,146.6 | 573.8 | 1,129.6 | 1,128.6 | 10 |
| 2 | 229.1 | | | 211.1 | L | 1,031.6 | 516.3 | 1,014.5 | 1,013.6 | 9 |
| 3 | 300.2 | | | 282.1 | A | 918.5 | 459.7 | 901.5 | 900.5 | 8 |
| 4 | 429.2 | | | 411.2 | E | 847.5 | 424.2 | 830.4 | 829.4 | 7 |
| 5 | 542.3 | | | 524.3 | I | 718.4 | 359.7 | 701.4 | 700.4 | 6 |
| 6 | 643.3 | 322.2 | | 625.3 | T | 605.3 | | 588.3 | 587.3 | 5 |
| 7 | 744.4 | 372.7 | | 726.4 | T | 504.3 | | 487.3 | 486.3 | 4 |
| 8 | 857.5 | 429.2 | | 839.5 | L | 403.2 | | 386.2 | 385.2 | 3 |
| 9 | 972.5 | 486.7 | | 954.5 | D | 290.1 | | 273.1 | 272.1 | 2 |
| 10 | 1,146.6 | 573.8 | 1,129.6 | 1,128.6 | R | 175.1 | | 158.1 | | 1 |

Unique Peptide Search: ERp44 (one peptide sequence; two peptides)

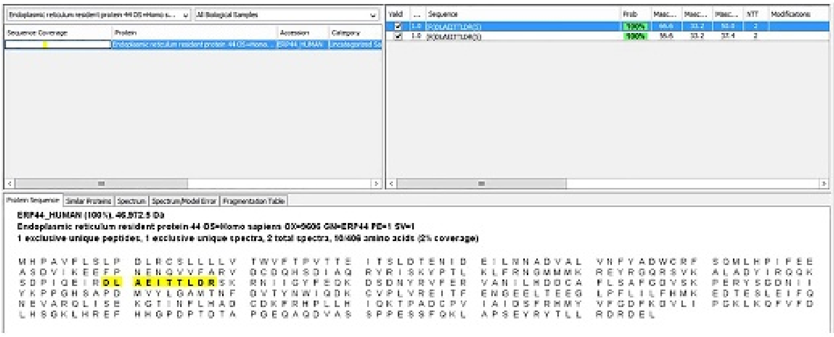

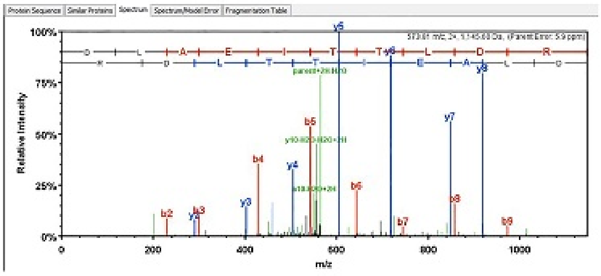

| B | B Ions | B+2H | B-NH3 | B-H2O | AA | Y Ions | Y+2H | Y-NH3 | Y-H2O | Y |
|---|---|---|---|---|---|---|---|---|---|---|
| 1 | 116.0 | | | 98.0 | D | 1,146.6 | 573.8 | 1,129.6 | 1,128.6 | 10 |
| 2 | 229.1 | | | 211.1 | L | 1,031.6 | 516.3 | 1,014.5 | 1,013.6 | 9 |
| 3 | 300.2 | | | 282.1 | A | 918.5 | 459.7 | 901.5 | 900.5 | 8 |
| 4 | 429.2 | | | 411.2 | E | 847.5 | 424.2 | 830.4 | 829.4 | 7 |
| 5 | 542.3 | | | 524.3 | I | 718.4 | 359.7 | 701.4 | 700.4 | 6 |
| 6 | 643.3 | 322.2 | | 625.3 | T | 605.3 | | 588.3 | 587.3 | 5 |
| 7 | 744.4 | 372.7 | | 726.4 | T | 504.3 | | 487.3 | 486.3 | 4 |
| 8 | 857.5 | 429.2 | | 839.5 | L | 403.2 | | 386.2 | 385.2 | 3 |
| 9 | 972.5 | 486.7 | | 954.5 | D | 290.1 | | 273.1 | 272.1 | 2 |
| 10 | 1,146.6 | 573.8 | 1,129.6 | 1,128.6 | R | 175.1 | | 158.1 | | 1 |

# S2B

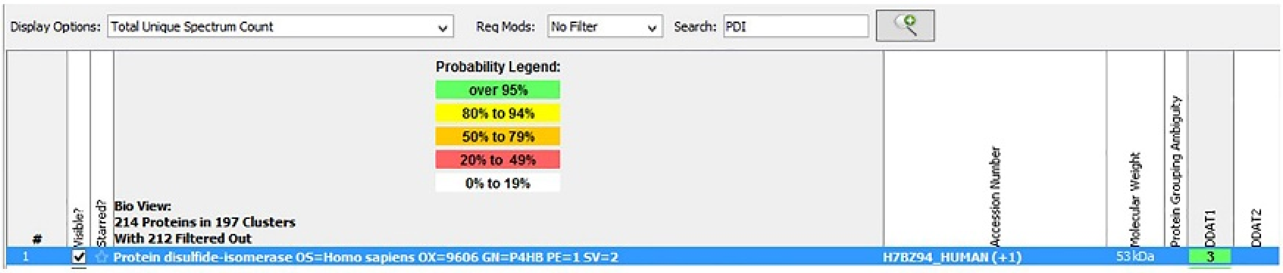


Unique Peptide Search: PDIs (three peptides sequences; 13 peptides)

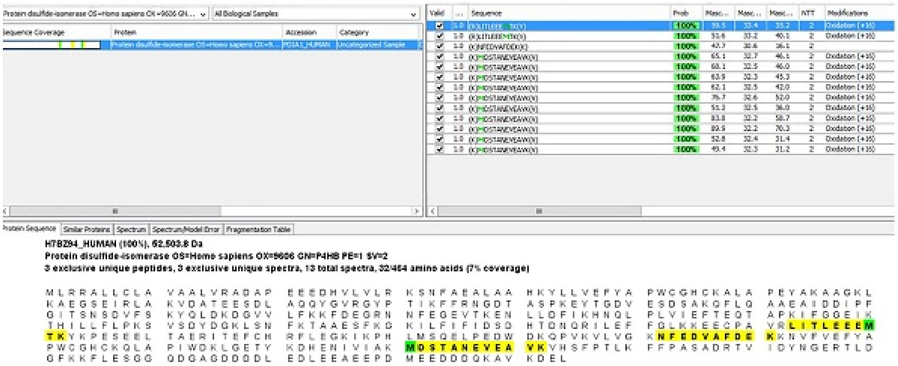


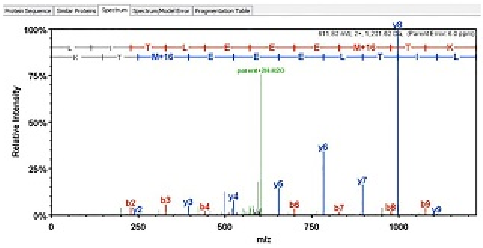


| B | B Ions | B+2H | B-NH3 | B-H2O | AA | Y Ions | Y+2H | Y-NH3 | Y-H2O | Y |
|---|---|---|---|---|---|---|---|---|---|---|
| 1 | 114.1 | | | | L | 1,222.6 | 611.8 | 1,205.6 | 1,204.6 | 10 |
| 2 | 227.2 | | | | I | 1,109.5 | 555.3 | 1,092.5 | 1,091.5 | 9 |
| 3 | 328.2 | | | 310.2 | T | 996.5 | 498.7 | 979.4 | 978.4 | 8 |
| 4 | 441.3 | | | 423.3 | L | 895.4 | 448.2 | 878.4 | 877.4 | 7 |
| 5 | 570.3 | | | 552.3 | E | 782.3 | 391.7 | 765.3 | 764.3 | 6 |
| 6 | 699.4 | 350.2 | | 681.4 | E | 653.3 | | 636.3 | 635.3 | 5 |
| 7 | 828.4 | 414.7 | | 810.4 | E | 524.2 | | 507.2 | 506.2 | 4 |
| 8 | 975.5 | 488.2 | | 957.5 | M+16 | 395.2 | | 378.2 | 377.2 | 3 |
| 9 | 1,076.5 | 538.8 | | 1,058.5 | T | 248.2 | | 231.1 | 230.1 | 2 |
| 10 | 1,222.6 | 611.8 | 1,205.6 | 1,204.6 | K | 147.1 | | 130.1 | | 1 |

Unique Peptide Search: PDIs (three peptide sequences; 13 peptides)

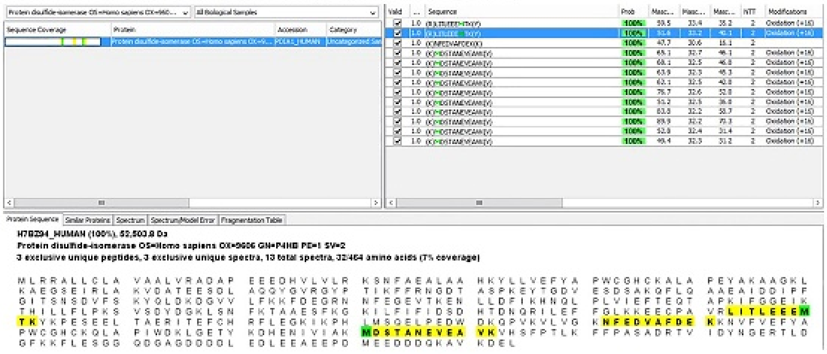


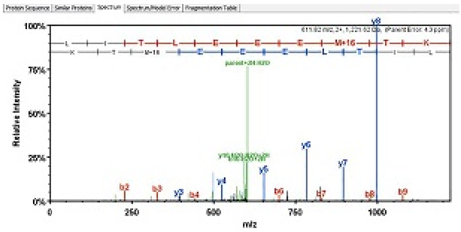


| B | B Ions | B+2H | B-NH3 | B-H2O | AA | Y Ions | Y+2H | Y-NH3 | Y-H2O | Y |
|---|---|---|---|---|---|---|---|---|---|---|
| 1 | 114.1 | | | | L | 1,222.6 | 611.8 | 1,205.6 | 1,204.6 | 10 |
| 2 | 227.2 | | | | I | 1,109.5 | 555.3 | 1,092.5 | 1,091.5 | 9 |
| 3 | 328.2 | | | 310.2 | T | 996.5 | 498.7 | 979.4 | 978.4 | 8 |
| 4 | 441.3 | | | 423.3 | L | 895.4 | 448.2 | 878.4 | 877.4 | 7 |
| 5 | 570.3 | | | 552.3 | E | 782.3 | 391.7 | 765.3 | 764.3 | 6 |
| 6 | 699.4 | 350.2 | | 681.4 | E | 653.3 | | 636.3 | 635.3 | 5 |
| 7 | 828.4 | 414.7 | | 810.4 | E | 524.2 | | 507.2 | 506.2 | 4 |
| 8 | 975.5 | 488.2 | | 957.5 | M+16 | 395.2 | | 378.2 | 377.2 | 3 |
| 9 | 1,076.5 | 538.8 | | 1,058.5 | T | 248.2 | | 231.1 | 230.1 | 2 |
| 10 | 1,222.6 | 611.8 | 1,205.6 | 1,204.6 | K | 147.1 | | 130.1 | | 1 |

### Unique Peptide Search: PDIs (three peptide sequences; 13 peptides)

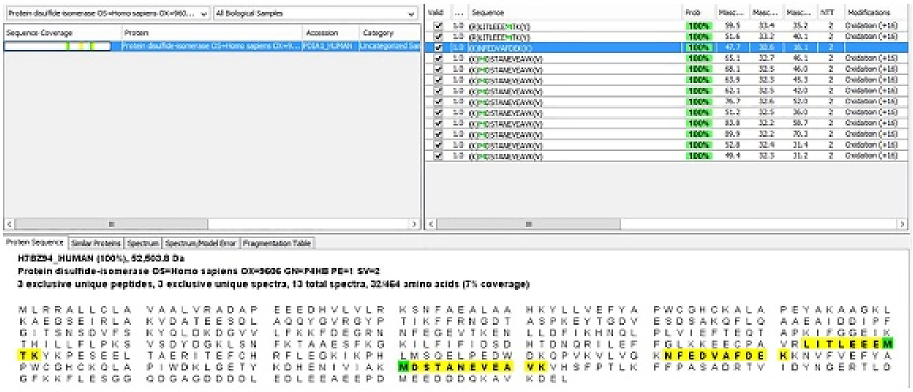


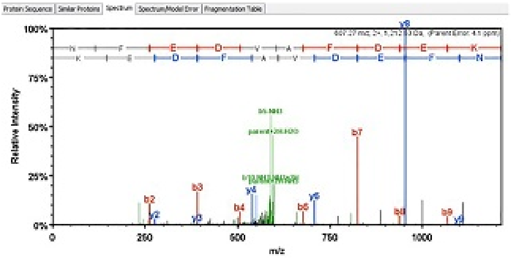

| B | B Ions | B+2H | B-NH3 | B-H2O | AA | Y Ions | Y+2H | Y-NH3 | Y-H2O | Y |
|---|---|---|---|---|---|---|---|---|---|---|
| 1 | 115.1 | | 98.0 | | N | 1,213.5 | 607.3 | 1,196.5 | 1,195.5 | 10 |
| 2 | 262.1 | | 245.1 | | F | 1,099.5 | 550.3 | 1,082.5 | 1,081.5 | 9 |
| 3 | 391.2 | | 374.1 | 373.2 | E | 952.4 | 476.7 | 935.4 | 934.4 | 8 |
| 4 | 506.2 | | 489.2 | 488.2 | D | 823.4 | 412.2 | 806.4 | 805.4 | 7 |
| 5 | 605.3 | | 588.2 | 587.2 | V | 708.4 | 354.7 | 691.3 | 690.3 | 6 |
| 6 | 676.3 | 338.7 | 659.3 | 658.3 | A | 609.3 | | 592.3 | 591.3 | 5 |
| 7 | 823.4 | 412.2 | 806.3 | 805.4 | F | 538.3 | | 521.2 | 520.2 | 4 |
| 8 | 938.4 | 469.7 | 921.4 | 920.4 | D | 391.2 | | 374.2 | 373.2 | 3 |
| 9 | 1,067.4 | 534.2 | 1,050.4 | 1,049.4 | E | 276.2 | | 259.1 | 258.1 | 2 |
| 10 | 1,213.5 | 607.3 | 1,196.5 | 1,195.5 | K | 147.1 | | 130.1 | | 1 |

### Unique Peptide Search: PDIs (three peptide sequences; 13 peptides)

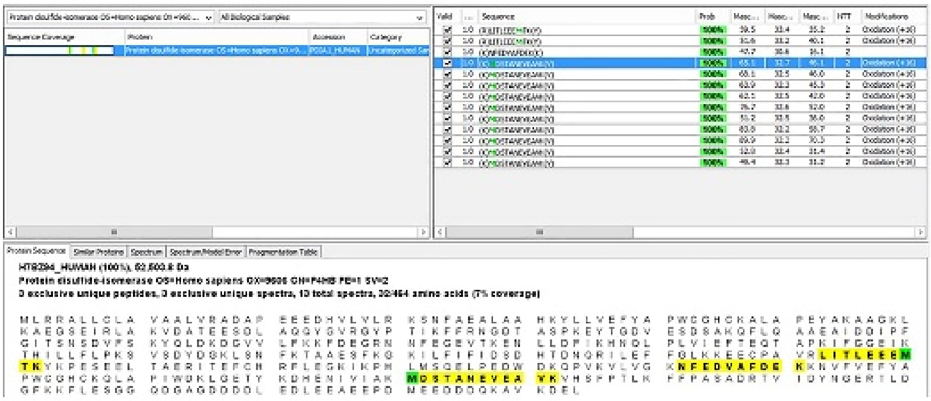


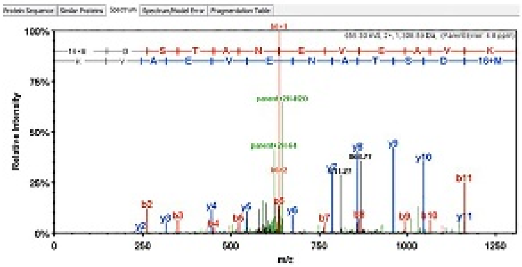

| B | B Ions | B+2H | B-NH3 | B-H2O | AA | Y Ions | Y+2H | Y-NH3 | Y-H2O | Y |
|---|---|---|---|---|---|---|---|---|---|---|
| 1 | 148.0 | | | | M+16 | 1,309.6 | 655.3 | 1,292.6 | 1,291.6 | 12 |
| 2 | 263.1 | | | 245.1 | D | 1,162.6 | 581.8 | 1,145.5 | 1,144.5 | 11 |
| 3 | 350.1 | | | 332.1 | S | 1,047.5 | 524.3 | 1,030.5 | 1,029.5 | 10 |
| 4 | 451.1 | | | 433.1 | T | 960.5 | 480.8 | 943.5 | 942.5 | 9 |
| 5 | 522.2 | | | 504.2 | A | 859.5 | 430.2 | 842.4 | 841.4 | 8 |
| 6 | 636.2 | 318.6 | 619.2 | 618.2 | N | 788.4 | 394.7 | 771.4 | 770.4 | 7 |
| 7 | 765.3 | 383.1 | 748.2 | 747.3 | E | 674.4 | 337.7 | 657.3 | 656.4 | 6 |
| 8 | 864.3 | 432.7 | 847.3 | 846.3 | V | 545.3 | | 528.3 | 527.3 | 5 |
| 9 | 993.4 | 497.2 | 976.4 | 975.4 | E | 446.3 | | 429.2 | 428.3 | 4 |
| 10 | 1,064.4 | 532.7 | 1,047.4 | 1,046.4 | A | 317.2 | | 300.2 | | 3 |
| 11 | 1,163.5 | 582.2 | 1,146.5 | 1,145.5 | V | 246.2 | | 229.2 | | 2 |
| 12 | 1,309.6 | 655.3 | 1,292.6 | 1,291.6 | K | 147.1 | | 130.1 | | 1 |

### Unique Peptide Search: PDIs (three peptide sequences; 13 peptides)

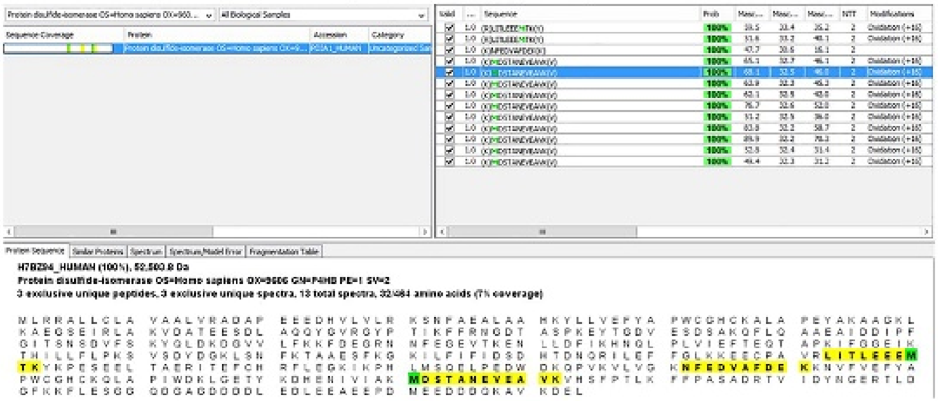


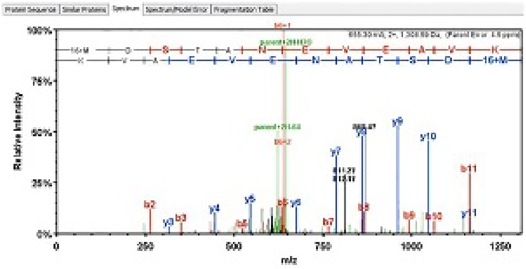


| B | B Ions | B+2H | B-NH3 | B-H2O | AA | Y Ions | Y+2H | Y-NH3 | Y-H2O | Y |
|---|---|---|---|---|---|---|---|---|---|---|
| 1 | 148.0 | | | | M+16 | 1,309.6 | 655.3 | 1,292.6 | 1,291.6 | 12 |
| 2 | 263.1 | | | 245.1 | D | 1,162.6 | 581.8 | 1,145.5 | 1,144.5 | 11 |
| 3 | 350.1 | | | 332.1 | S | 1,047.5 | 524.3 | 1,030.5 | 1,029.5 | 10 |
| 4 | 451.1 | | | 433.1 | T | 960.5 | 480.8 | 943.5 | 942.5 | 9 |
| 5 | 522.2 | | | 504.2 | A | 859.5 | 430.2 | 842.4 | 841.4 | 8 |
| 6 | 636.2 | 318.6 | 619.2 | 618.2 | N | 788.4 | 394.7 | 771.4 | 770.4 | 7 |
| 7 | 765.3 | 383.1 | 748.2 | 747.3 | E | 674.4 | 337.7 | 657.3 | 656.4 | 6 |
| 8 | 864.3 | 432.7 | 847.3 | 846.3 | V | 545.3 | | 528.3 | 527.3 | 5 |
| 9 | 993.4 | 497.2 | 976.4 | 975.4 | E | 446.3 | | 429.2 | 428.3 | 4 |
| 10 | 1,064.4 | 532.7 | 1,047.4 | 1,046.4 | A | 317.2 | | 300.2 | | 3 |
| 11 | 1,163.5 | 582.2 | 1,146.5 | 1,145.5 | V | 246.2 | | 229.2 | | 2 |
| 12 | 1,309.6 | 655.3 | 1,292.6 | 1,291.6 | K | 147.1 | | 130.1 | | 1 |

### Unique Peptide Search: PDIs (three peptide sequences; 13 peptides)

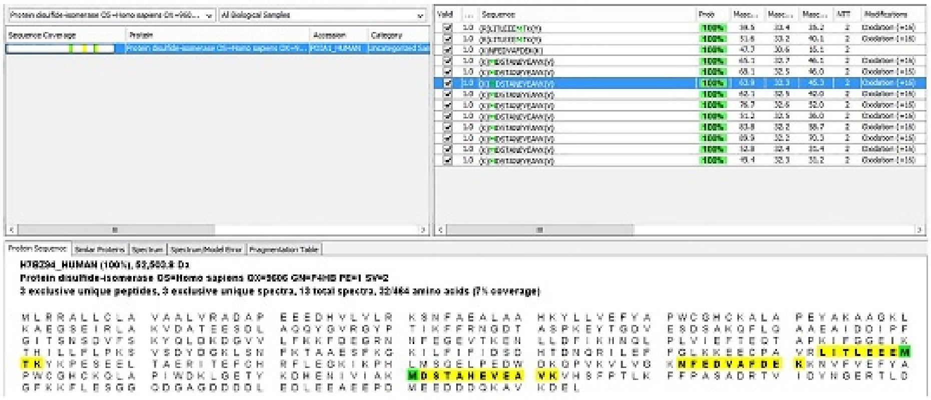


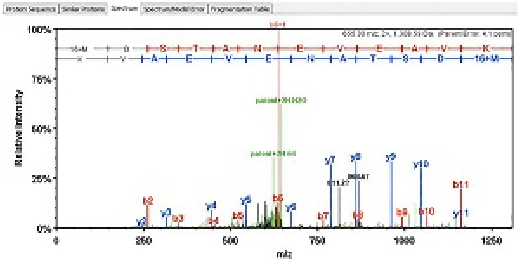


| B | B Ions | B+2H | B-NH3 | B-H2O | AA | Y Ions | Y+2H | Y-NH3 | Y-H2O | Y |
|---|---|---|---|---|---|---|---|---|---|---|
| 1 | 148.0 | | | | M+16 | 1,309.6 | 655.3 | 1,292.6 | 1,291.6 | 12 |
| 2 | 263.1 | | | 245.1 | D | 1,162.6 | 581.8 | 1,145.5 | 1,144.5 | 11 |
| 3 | 350.1 | | | 332.1 | S | 1,047.5 | 524.3 | 1,030.5 | 1,029.5 | 10 |
| 4 | 451.1 | | | 433.1 | T | 960.5 | 480.8 | 943.5 | 942.5 | 9 |
| 5 | 522.2 | | | 504.2 | A | 859.5 | 430.2 | 842.4 | 841.4 | 8 |
| 6 | 636.2 | 318.6 | 619.2 | 618.2 | N | 788.4 | 394.7 | 771.4 | 770.4 | 7 |
| 7 | 765.3 | 383.1 | 748.2 | 747.3 | E | 674.4 | 337.7 | 657.3 | 656.4 | 6 |
| 8 | 864.3 | 432.7 | 847.3 | 846.3 | V | 545.3 | | 528.3 | 527.3 | 5 |
| 9 | 993.4 | 497.2 | 976.4 | 975.4 | E | 446.3 | | 429.2 | 428.3 | 4 |
| 10 | 1,064.4 | 532.7 | 1,047.4 | 1,046.4 | A | 317.2 | | 300.2 | | 3 |
| 11 | 1,163.5 | 582.2 | 1,146.5 | 1,145.5 | V | 246.2 | | 229.2 | | 2 |
| 12 | 1,309.6 | 655.3 | 1,292.6 | 1,291.6 | K | 147.1 | | 130.1 | | 1 |

Unique Peptide Search: PDIs (three peptide sequences; 13 peptides)

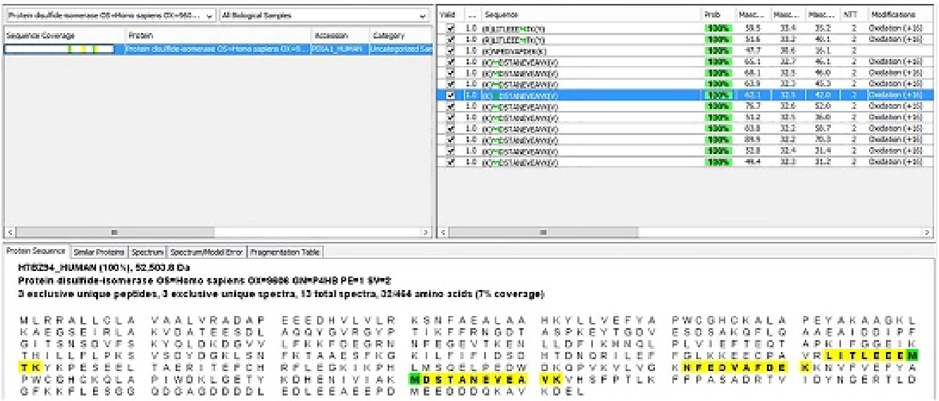

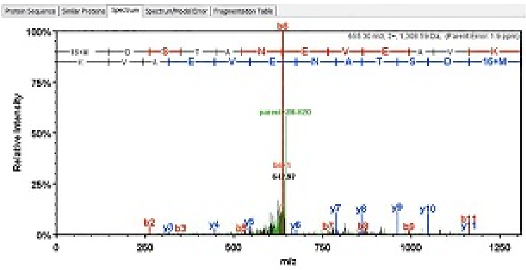

| B | B Ions | B+2H | B-NH3 | B-H2O | AA | Y Ions | Y+2H | Y-NH3 | Y-H2O | Y |
|---|---|---|---|---|---|---|---|---|---|---|
| 1 | 148.0 | | | | M+16 | 1,309.6 | 655.3 | 1,292.6 | 1,291.6 | 12 |
| 2 | 263.1 | | | 245.1 | D | 1,162.6 | 581.8 | 1,145.5 | 1,144.5 | 11 |
| 3 | 350.1 | | | 332.1 | S | 1,047.5 | 524.3 | 1,030.5 | 1,029.5 | 10 |
| 4 | 451.1 | | | 433.1 | T | 960.5 | 480.8 | 943.5 | 942.5 | 9 |
| 5 | 522.2 | | | 504.2 | A | 859.5 | 430.2 | 842.4 | 841.4 | 8 |
| 6 | 636.2 | 318.6 | 619.2 | 618.2 | N | 788.4 | 394.7 | 771.4 | 770.4 | 7 |
| 7 | 765.3 | 383.1 | 748.2 | 747.3 | E | 674.4 | 337.7 | 657.3 | 656.4 | 6 |
| 8 | 864.3 | 432.7 | 847.3 | 846.3 | V | 545.3 | | 528.3 | 527.3 | 5 |
| 9 | 993.4 | 497.2 | 976.4 | 975.4 | E | 446.3 | | 429.2 | 428.3 | 4 |
| 10 | 1,064.4 | 532.7 | 1,047.4 | 1,046.4 | A | 317.2 | | 300.2 | | 3 |
| 11 | 1,163.5 | 582.2 | 1,146.5 | 1,145.5 | V | 246.2 | | 229.2 | | 2 |
| 12 | 1,309.6 | 655.3 | 1,292.6 | 1,291.6 | K | 147.1 | | 130.1 | | 1 |

Unique Peptide Search: PDIs (three peptide sequences; 13 peptides)

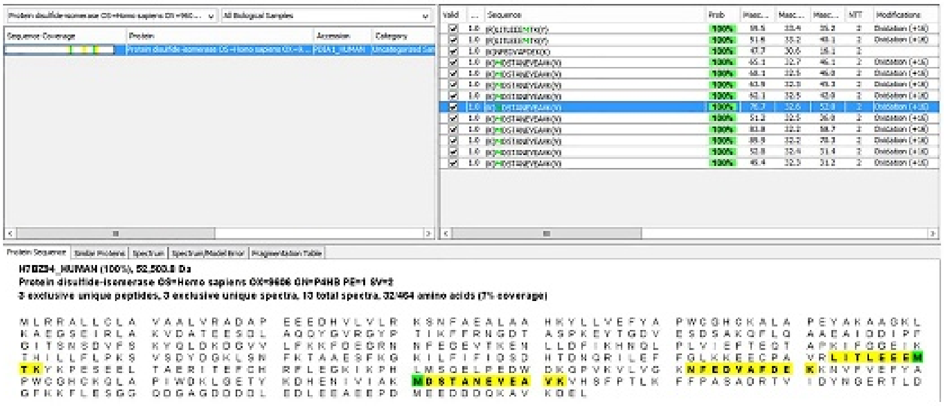

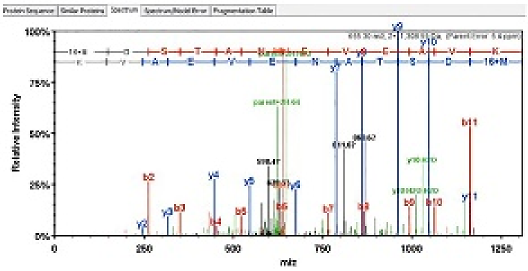

| B | B Ions | B+2H | B-NH3 | B-H2O | AA | Y Ions | Y+2H | Y-NH3 | Y-H2O | Y |
|---|---|---|---|---|---|---|---|---|---|---|
| 1 | 148.0 | | | | M+16 | 1,309.6 | 655.3 | 1,292.6 | 1,291.6 | 12 |
| 2 | 263.1 | | | 245.1 | D | 1,162.6 | 581.8 | 1,145.5 | 1,144.5 | 11 |
| 3 | 350.1 | | | 332.1 | S | 1,047.5 | 524.3 | 1,030.5 | 1,029.5 | 10 |
| 4 | 451.1 | | | 433.1 | T | 960.5 | 480.8 | 943.5 | 942.5 | 9 |
| 5 | 522.2 | | | 504.2 | A | 859.5 | 430.2 | 842.4 | 841.4 | 8 |
| 6 | 636.2 | 318.6 | 619.2 | 618.2 | N | 788.4 | 394.7 | 771.4 | 770.4 | 7 |
| 7 | 765.3 | 383.1 | 748.2 | 747.3 | E | 674.4 | 337.7 | 657.3 | 656.4 | 6 |
| 8 | 864.3 | 432.7 | 847.3 | 846.3 | V | 545.3 | | 528.3 | 527.3 | 5 |
| 9 | 993.4 | 497.2 | 976.4 | 975.4 | E | 446.3 | | 429.2 | 428.3 | 4 |
| 10 | 1,064.4 | 532.7 | 1,047.4 | 1,046.4 | A | 317.2 | | 300.2 | | 3 |
| 11 | 1,163.5 | 582.2 | 1,146.5 | 1,145.5 | V | 246.2 | | 229.2 | | 2 |
| 12 | 1,309.6 | 655.3 | 1,292.6 | 1,291.6 | K | 147.1 | | 130.1 | | 1 |

### Unique Peptide Search: PDIs (three peptide sequences; 13 peptides)

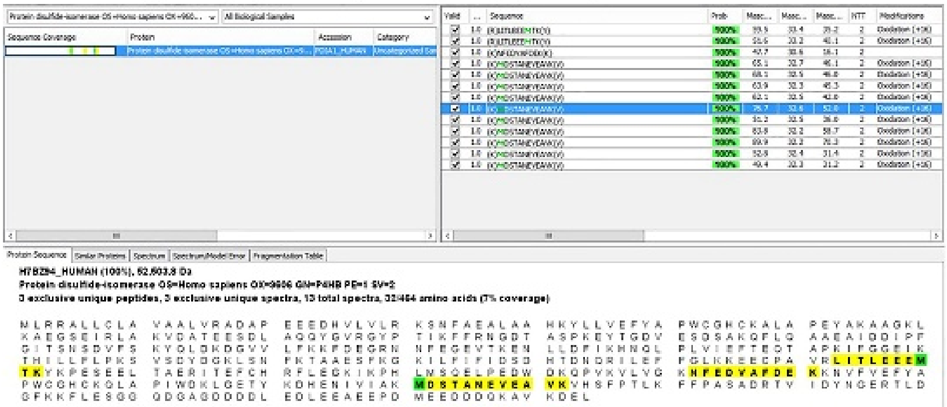


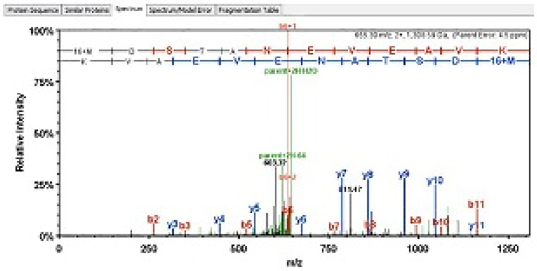


| B | B Ions | B+2H | B-NH3 | B-H2O | AA | Y Ions | Y+2H | Y-NH3 | Y-H2O | Y |
|---|---|---|---|---|---|---|---|---|---|---|
| 1 | 148.0 | | | | M+16 | 1,309.6 | 655.3 | 1,292.6 | 1,291.6 | 12 |
| 2 | 263.1 | | | 245.1 | D | 1,162.6 | 581.8 | 1,145.5 | 1,144.5 | 11 |
| 3 | 350.1 | | | 332.1 | S | 1,047.5 | 524.3 | 1,030.5 | 1,029.5 | 10 |
| 4 | 451.1 | | | 433.1 | T | 960.5 | 480.8 | 943.5 | 942.5 | 9 |
| 5 | 522.2 | | | 504.2 | A | 859.5 | 430.2 | 842.4 | 841.4 | 8 |
| 6 | 636.2 | 318.6 | 619.2 | 618.2 | N | 788.4 | 394.7 | 771.4 | 770.4 | 7 |
| 7 | 765.3 | 383.1 | 748.2 | 747.3 | E | 674.4 | 337.7 | 657.3 | 656.4 | 6 |
| 8 | 864.3 | 432.7 | 847.3 | 846.3 | V | 545.3 | | 528.3 | 527.3 | 5 |
| 9 | 993.4 | 497.2 | 976.4 | 975.4 | E | 446.3 | | 429.2 | 428.3 | 4 |
| 10 | 1,064.4 | 532.7 | 1,047.4 | 1,046.4 | A | 317.2 | | 300.2 | | 3 |
| 11 | 1,163.5 | 582.2 | 1,146.5 | 1,145.5 | V | 246.2 | | 229.2 | | 2 |
| 12 | 1,309.6 | 655.3 | 1,292.6 | 1,291.6 | K | 147.1 | | 130.1 | | 1 |

### Unique Peptide Search: PDIs (three peptide sequences; 13 peptides)

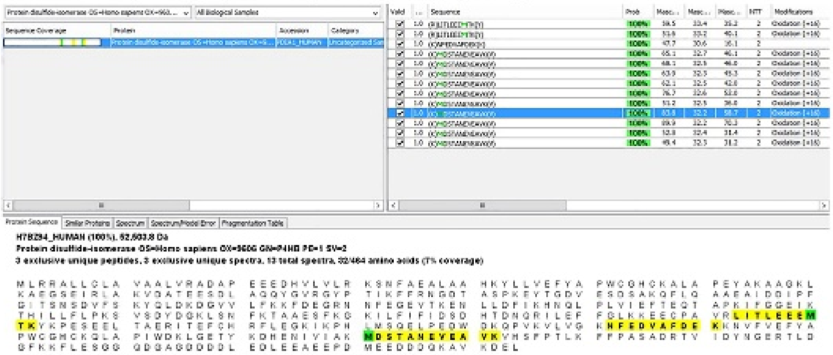


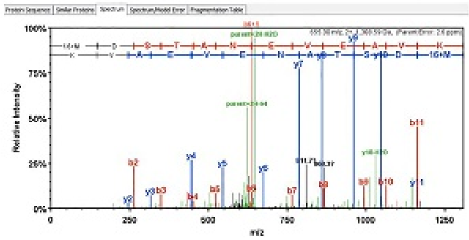


| B | B Ions | B+2H | B-NH3 | B-H2O | AA | Y Ions | Y+2H | Y-NH3 | Y-H2O | Y |
|---|---|---|---|---|---|---|---|---|---|---|
| 1 | 148.0 | | | | M+16 | 1,309.6 | 655.3 | 1,292.6 | 1,291.6 | 12 |
| 2 | 263.1 | | | 245.1 | D | 1,162.6 | 581.8 | 1,145.5 | 1,144.5 | 11 |
| 3 | 350.1 | | | 332.1 | S | 1,047.5 | 524.3 | 1,030.5 | 1,029.5 | 10 |
| 4 | 451.1 | | | 433.1 | T | 960.5 | 480.8 | 943.5 | 942.5 | 9 |
| 5 | 522.2 | | | 504.2 | A | 859.5 | 430.2 | 842.4 | 841.4 | 8 |
| 6 | 636.2 | 318.6 | 619.2 | 618.2 | N | 788.4 | 394.7 | 771.4 | 770.4 | 7 |
| 7 | 765.3 | 383.1 | 748.2 | 747.3 | E | 674.4 | 337.7 | 657.3 | 656.4 | 6 |
| 8 | 864.3 | 432.7 | 847.3 | 846.3 | V | 545.3 | | 528.3 | 527.3 | 5 |
| 9 | 993.4 | 497.2 | 976.4 | 975.4 | E | 446.3 | | 429.2 | 428.3 | 4 |
| 10 | 1,064.4 | 532.7 | 1,047.4 | 1,046.4 | A | 317.2 | | 300.2 | | 3 |
| 11 | 1,163.5 | 582.2 | 1,146.5 | 1,145.5 | V | 246.2 | | 229.2 | | 2 |
| 12 | 1,309.6 | 655.3 | 1,292.6 | 1,291.6 | K | 147.1 | | 130.1 | | 1 |

### Unique Peptide Search: PDIs (three peptide sequences; 13 peptides)

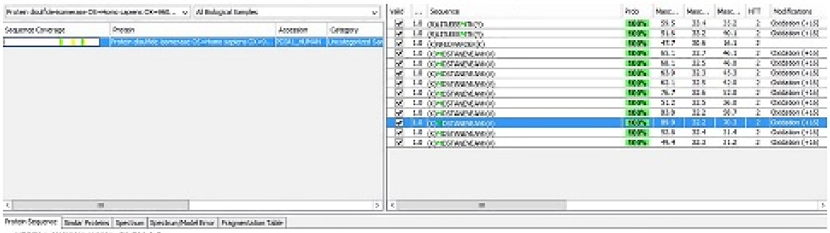

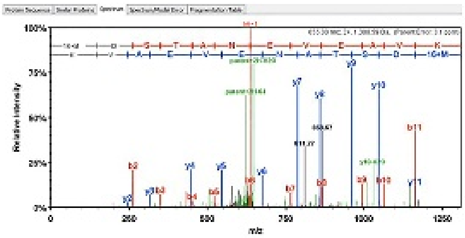

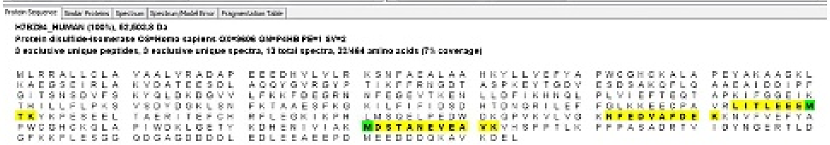

| B | B Ions | B+2H | B-NH3 | B-H2O | AA | Y Ions | Y+2H | Y-NH3 | Y-H2O | Y |
|---|---|---|---|---|---|---|---|---|---|---|
| 1 | 148.0 | | | | M+16 | 1,309.6 | 655.3 | 1,292.6 | 1,291.6 | 12 |
| 2 | 263.1 | | | 245.1 | D | 1,162.6 | 581.8 | 1,145.5 | 1,144.5 | 11 |
| 3 | 350.1 | | | 332.1 | S | 1,047.5 | 524.3 | 1,030.5 | 1,029.5 | 10 |
| 4 | 451.1 | | | 433.1 | T | 960.5 | 480.8 | 943.5 | 942.5 | 9 |
| 5 | 522.2 | | | 504.2 | A | 859.5 | 430.2 | 842.4 | 841.4 | 8 |
| 6 | 636.2 | 318.6 | 619.2 | 618.2 | N | 788.4 | 394.7 | 771.4 | 770.4 | 7 |
| 7 | 765.3 | 383.1 | 748.2 | 747.3 | E | 674.4 | 337.7 | 657.3 | 656.4 | 6 |
| 8 | 864.3 | 432.7 | 847.3 | 846.3 | V | 545.3 | | 528.3 | 527.3 | 5 |
| 9 | 993.4 | 497.2 | 976.4 | 975.4 | E | 446.3 | | 429.2 | 428.3 | 4 |
| 10 | 1,064.4 | 532.7 | 1,047.4 | 1,046.4 | A | 317.2 | | 300.2 | | 3 |
| 11 | 1,163.5 | 582.2 | 1,146.5 | 1,145.5 | V | 246.2 | | 229.2 | | 2 |
| 12 | 1,309.6 | 655.3 | 1,292.6 | 1,291.6 | K | 147.1 | | 130.1 | | 1 |

### Unique Peptide Search: PDIs (three peptide sequences; 13 peptides)

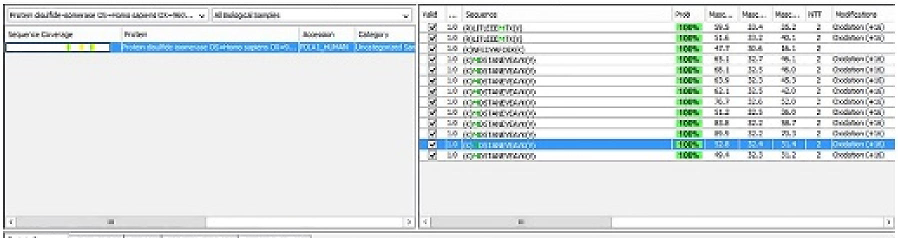

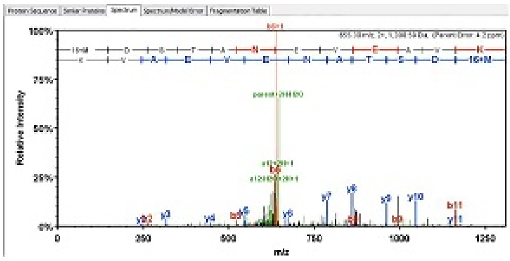

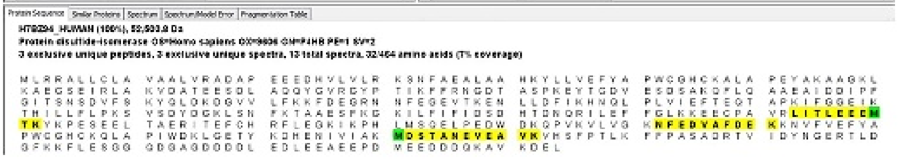

| B | B Ions | B+2H | B-NH3 | B-H2O | AA | Y Ions | Y+2H | Y-NH3 | Y-H2O | Y |
|---|---|---|---|---|---|---|---|---|---|---|
| 1 | 148.0 | | | | M+16 | 1,309.6 | 655.3 | 1,292.6 | 1,291.6 | 12 |
| 2 | 263.1 | | | 245.1 | D | 1,162.6 | 581.8 | 1,145.5 | 1,144.5 | 11 |
| 3 | 350.1 | | | 332.1 | S | 1,047.5 | 524.3 | 1,030.5 | 1,029.5 | 10 |
| 4 | 451.1 | | | 433.1 | T | 960.5 | 480.8 | 943.5 | 942.5 | 9 |
| 5 | 522.2 | | | 504.2 | A | 859.5 | 430.2 | 842.4 | 841.4 | 8 |
| 6 | 636.2 | 318.6 | 619.2 | 618.2 | N | 788.4 | 394.7 | 771.4 | 770.4 | 7 |
| 7 | 765.3 | 383.1 | 748.2 | 747.3 | E | 674.4 | 337.7 | 657.3 | 656.4 | 6 |
| 8 | 864.3 | 432.7 | 847.3 | 846.3 | V | 545.3 | | 528.3 | 527.3 | 5 |
| 9 | 993.4 | 497.2 | 976.4 | 975.4 | E | 446.3 | | 429.2 | 428.3 | 4 |
| 10 | 1,064.4 | 532.7 | 1,047.4 | 1,046.4 | A | 317.2 | | 300.2 | | 3 |
| 11 | 1,163.5 | 582.2 | 1,146.5 | 1,145.5 | V | 246.2 | | 229.2 | | 2 |
| 12 | 1,309.6 | 655.3 | 1,292.6 | 1,291.6 | K | 147.1 | | 130.1 | | 1 |

### Unique Peptide Search: PDIs (three peptide sequences; 13 peptides)

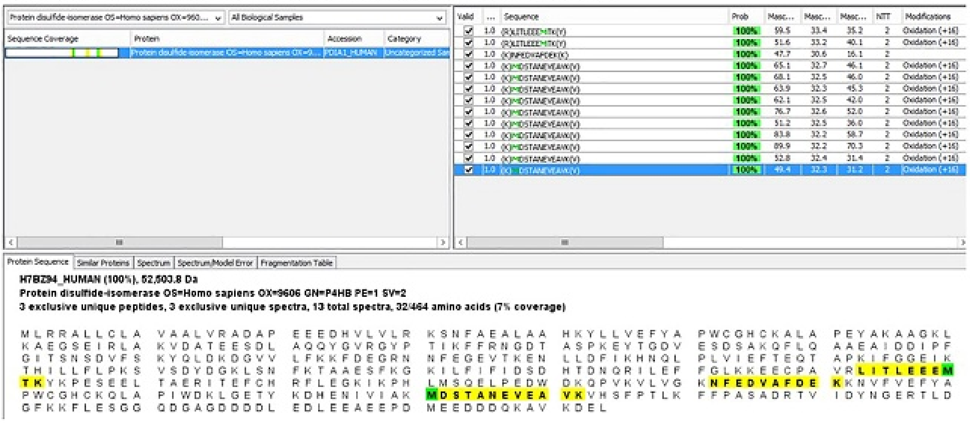


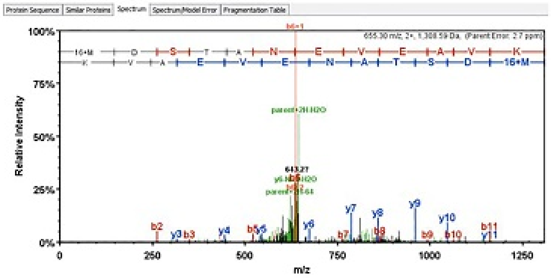


| B | B Ions | B+2H | B-NH3 | B-H2O | AA | Y Ions | Y+2H | Y-NH3 | Y-H2O | Y |
|---|---|---|---|---|---|---|---|---|---|---|
| 1 | 148.0 | | | | M+16 | 1,309.6 | 655.3 | 1,292.6 | 1,291.6 | 12 |
| 2 | 263.1 | | | 245.1 | D | 1,162.6 | 581.8 | 1,145.5 | 1,144.5 | 11 |
| 3 | 350.1 | | | 332.1 | S | 1,047.5 | 524.3 | 1,030.5 | 1,029.5 | 10 |
| 4 | 451.1 | | | 433.1 | T | 960.5 | 480.8 | 943.5 | 942.5 | 9 |
| 5 | 522.2 | | | 504.2 | A | 859.5 | 430.2 | 842.4 | 841.4 | 8 |
| 6 | 636.2 | 318.6 | 619.2 | 618.2 | N | 788.4 | 394.7 | 771.4 | 770.4 | 7 |
| 7 | 765.3 | 383.1 | 748.2 | 747.3 | E | 674.4 | 337.7 | 657.3 | 656.4 | 6 |
| 8 | 864.3 | 432.7 | 847.3 | 846.3 | V | 545.3 | | 528.3 | 527.3 | 5 |
| 9 | 993.4 | 497.2 | 976.4 | 975.4 | E | 446.3 | | 429.2 | 428.3 | 4 |
| 10 | 1,064.4 | 532.7 | 1,047.4 | 1,046.4 | A | 317.2 | | 300.2 | | 3 |
| 11 | 1,163.5 | 582.2 | 1,146.5 | 1,145.5 | V | 246.2 | | 229.2 | | 2 |
| 12 | 1,309.6 | 655.3 | 1,292.6 | 1,291.6 | K | 147.1 | | 130.1 | | 1 |

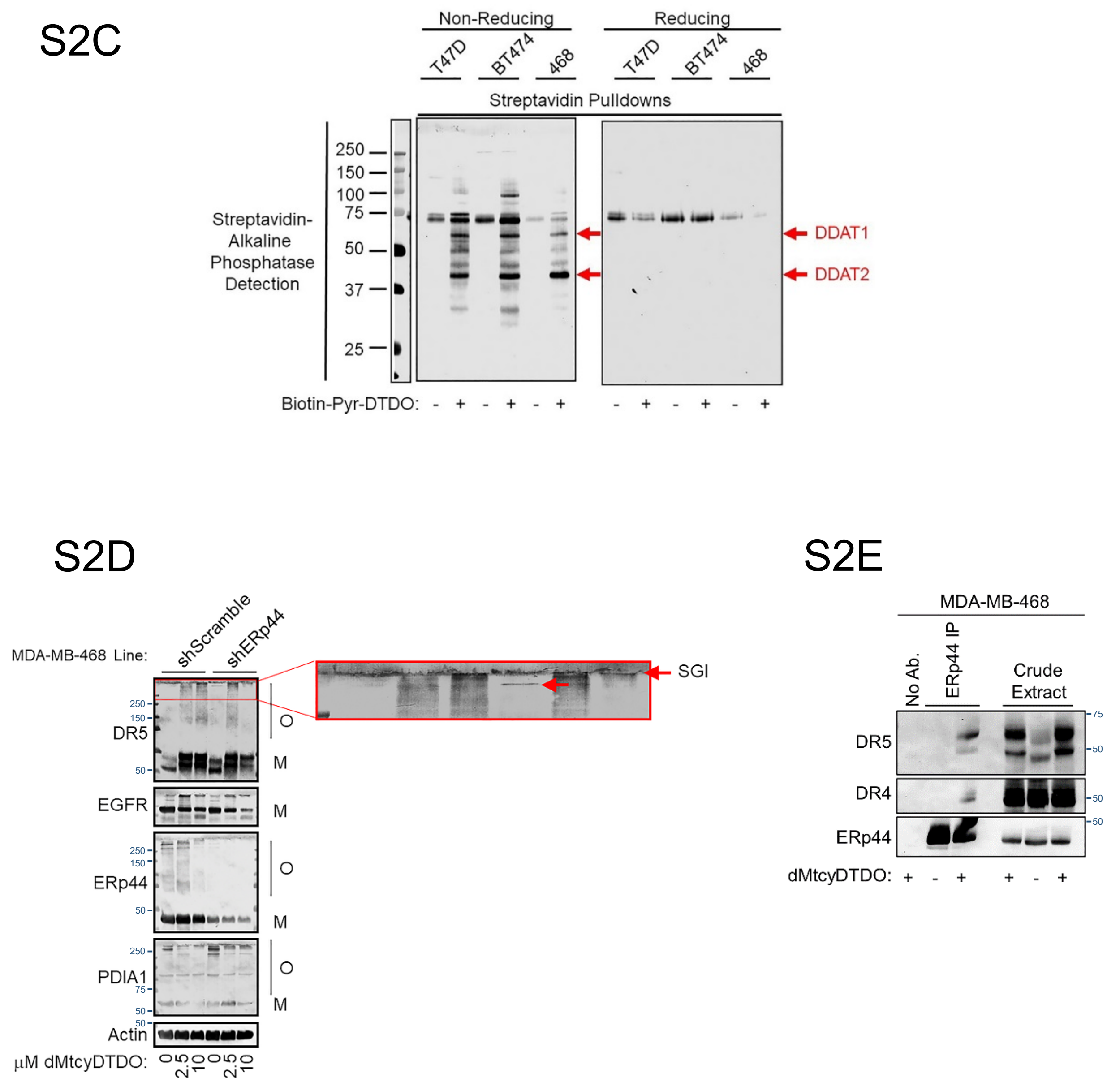


Fig. S2-Confirmation of ERp44 and PDIA1 as DDA targets. S2A. Peptide coverage of ERp44, and S2B. Peptide coverage of PDIA1 identified in Biotin-PyrDTDO/Streptavidin-Agarose pulldowns from MDA-MB-468 cells by tandem mass spectrometry. S2C. Streptavidin-Agarose pulldowns of Biotin-PyrDTDO treated cells performed in the presence of 100 mM NEM. Half of each sample was brought to 1 M 2-mercaptoethanol and boiled for 20 minutes to generate reduced samples. These non-reducing (left panel) or reducing samples (right panel) were analyzed by blotting with Streptavidin-Alkaline Phosphatase. Molecular weight markers are indicated in kiloDaltons. S2D. Biological replicate of the ERp44 knockdown experiment presented in Fig. 5C, D. Inset shows DR5 oligomerization upon ERp44 knockdown (red arrow). O and M represent oligomeric and monomeric protein forms. S2E. Biological replicate of the experiment presented in Fig. 7A.
